# Brain Region-Specific Epigenomic Reorganization and Altered Cell States in Alzheimer’s Disease

**DOI:** 10.1101/2025.09.29.678849

**Authors:** Wenliang Wang, Peter Berube, Bing Yang, Rosa Castanon, Anna Bartlett, Kaushik Komandur, Joseph R. Nery, Cesar Barragan, Mia Kenworthy, Cynthia Valadon, Jordan Altshul, Alaina Petrella, Derek Chan, Chumo Chen, Andrea Saldaña Acerbo, Jammy Luo, Manya Jain, Eshaan Soma, Huaming Chen, Michelle Liem, Mikayla Marrin, Caz O’Connor, Nathan Zemke, Derek Oakley, Bing Ren, Bradley T. Hyman, Joseph R. Ecker

## Abstract

Alzheimer’s disease (AD) is the most common neurodegenerative disorder, yet the molecular mechanisms underlying its region– and cell-type-specific pathogenesis remain poorly defined. Here, we generated a large-scale, single-cell multi-omic atlas—integrating DNA methylation and 3D genome architecture—from postmortem brain tissue of matched AD patients and cognitively normal controls. Samples were collected from three brain regions with distinct vulnerability to AD pathology: the temporal cortex (TC), primary visual cortex (VC), and prefrontal cortex (PFC). Our dataset comprises over 230,000 individual cells, spanning major neuronal and glial populations, and provides a high-resolution view of multi-layer epigenomic regulation. We identified widespread AD-associated DNA methylation changes and marked reorganization of 3D genome structure, including alterations in A/B compartments, topologically associating domains (TADs), and chromatin loops. These changes are strongly region-specific: TC displays pronounced hypermethylation, transcriptional downregulation, and elevated boundary density, whereas VC shows opposing trends and PFC an intermediate profile. We further uncovered previously unrecognized AD-associated glial and neuronal states defined by coordinated epigenomic dysregulation and recurrent genomic deletions, particularly near telomeric regions. This region-resolved, single-cell multi-omic atlas reveals divergent epigenomic trajectories across brain regions and cell types in AD, offering new mechanistic insights and a framework for targeted therapeutic strategies.

## Introduction

Alzheimer’s disease (AD) is the most prevalent neurodegenerative disorder, affecting over 50 million people worldwide and posing an increasing public health burden as populations age. Clinically, AD is characterized by progressive cognitive decline, while neuropathologically it is defined by extracellular amyloid-β plaques, intracellular neurofibrillary tangles composed of hyperphosphorylated tau, and widespread neuronal and synaptic loss^1,2^. Although genetic and transcriptomic studies have identified numerous AD-associated risk loci^3^ and expression signatures^4–9^, recent mechanistic investigations of selective vulnerability across brain regions and cell types have focused primarily on chromatin accessibility^10,11^, leaving other molecular determinants largely unexplored. Increasing evidence indicates that additional epigenomic modalities—such as DNA methylation and higher-order chromatin architecture—may critically mediate the interplay between genetic and environmental risk factors^12^, thereby contributing to the early initiation and progression of disease^13,14^.

Epigenetic dysregulation has emerged as a hallmark of Alzheimer’s disease (AD), particularly evident through widespread alterations in DNA methylation, histone modifications, chromatin accessibility, and 3D genome architecture—alterations initially revealed by bulk epigenomic studies that collectively impair neuronal gene regulation and cellular identity^15–18^. Epigenome-wide and single-cell studies have uncovered both locus-specific and global disruptions, especially in synaptic, immune, and metabolic networks^11^. A compelling recent development is the concept of “epigenome erosion,” a phenomenon first described in single-cell epigenomic analyses of late-stage AD, where chromatin accessibility landscapes progressively lose cell-type specificity, indicating loss of regulatory fidelity and neuronal identity^10,11^. However, to date, this erosion concept is primarily based on changes in chromatin accessibility, while the extent to which other epigenetic layers—such as DNA methylation and 3D chromosomal architecture—are similarly affected in AD remains largely unexplored.

To comprehensively map the regulatory landscape of Alzheimer’s disease at cellular resolution, we performed single-cell multi-omic profiling of postmortem brain tissue from AD patients and cognitively normal controls across three brain regions: temporal cortex (TC), primary visual cortex (VC), and prefrontal cortex (PFC). We jointly profiled DNA methylation and chromatin conformation across hundreds of thousands of cells, and in a companion study using the same samples, we also measured chromatin accessibility and gene expression. Our analysis revealed widespread and cell–type–specific alterations in DNA methylation and 3D genome architecture in AD. Importantly, we observed strong regional specificity: the temporal and visual cortices exhibited opposing AD-associated regulatory signatures, while the prefrontal cortex showed intermediate patterns. We identified disease-associated cell states shared across cell types, marked by coordinated changes across all molecular layers and enriched for genomic deletions proximal to telomeres. Together, our study presents a comprehensive, spatially resolved single-cell epigenomic atlas of AD, highlighting region– and cell–type–specific regulatory programs underlying disease pathology.

## Results

### Single-cell multi-omics profiling of Alzheimer’s brains

We investigated three anatomically and pathologically distinct brain regions—the primary visual cortex (VC), prefrontal cortex (PFC), and temporal cortex (TC)—from eleven individuals with Alzheimer’s disease (AD) and nine cognitively healthy controls (Supplementary Table 1). These regions span a gradient of AD vulnerability, with the VC being the least affected, PFC showing intermediate involvement, and the TC among the most severely impacted. From each sample, we performed single-nucleus methyl-3C sequencing (snm3C-seq)^19^, a joint assay combining chromosome conformation capture (3C) and DNA methylation (DNAm), and utilized the 10x Genomics Chromium Single Cell Multiome ATAC + Gene Expression (10x multiome) data from our companion study. After rigorous quality filtering, we sorted at least 3,000 cells for each sample (Extended Data Fig. 1A), and retained 221,245 cells for methylation and 3C analysis (Figure 1A, Extended Data Fig. 1B-D). While DNA methylation metrics consistently indicated high data quality, additional quality control criteria applied to chromatin conformation data reduced the 3C dataset to 191,151 high-quality cells (Extended Data Fig. 1C), yielding approximately 3,000 cells per donor per region for downstream analysis. Our primary analyses focused on DNA methylation and 3D genome architecture, leveraging the integrated methylome-conformation modality. A companion study provides an in-depth analysis of chromatin accessibility and gene expression from the same cohort. Together, these datasets provide a comprehensive multi-omic view of brain region and cell–type–specific epigenomic alterations in AD.

**Figure 1.**
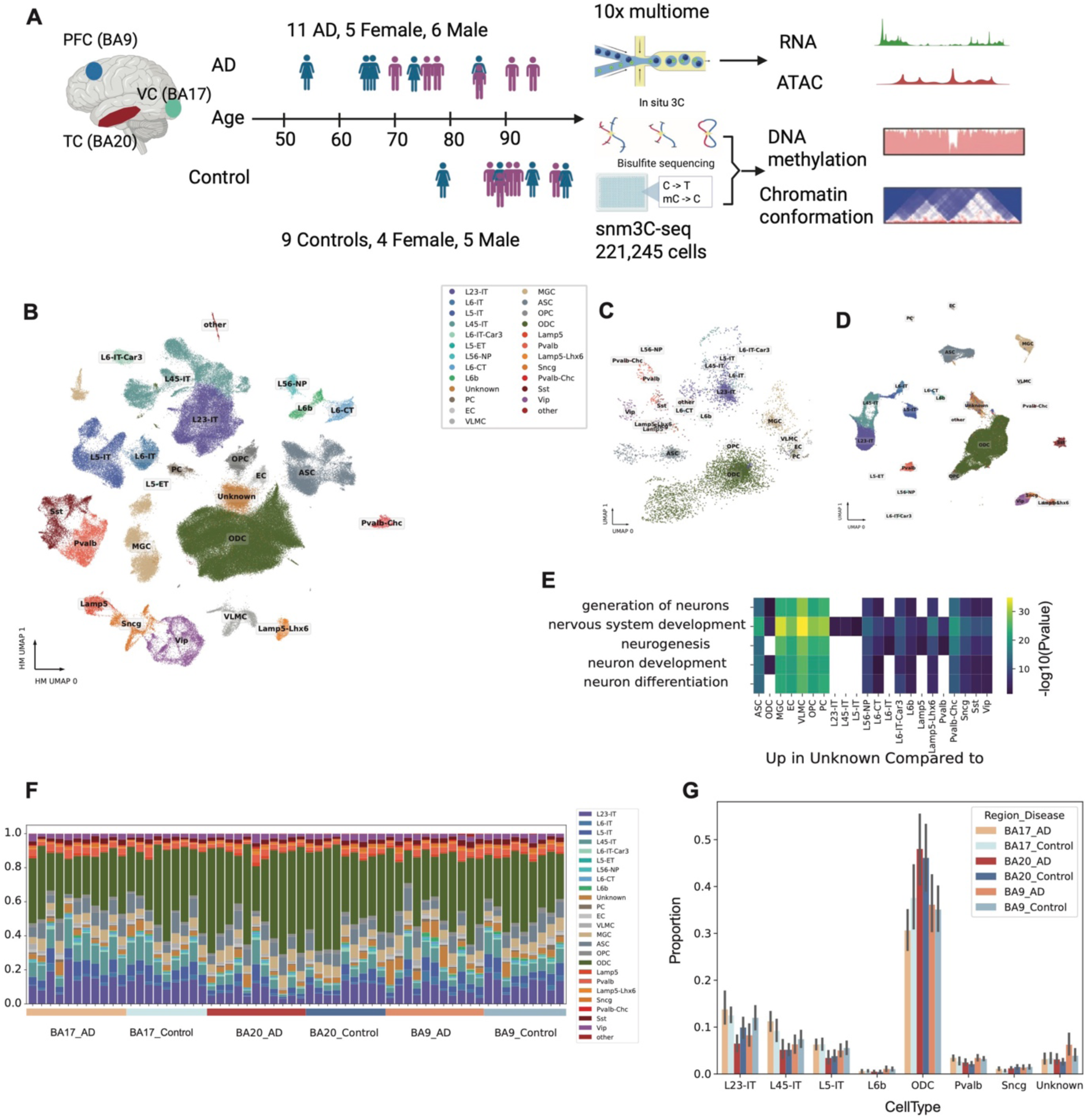
Single-cell multi-omic and multi-region profiling of Alzheimer’s disease (AD) and healthy control brains. (A) Overview of sample collection and single-cell multi-omic profiling in this study. (B) Uniform Manifold Approximation and Projection (UMAP) visualization after Harmony integration of 5 kb resolution CG DNA methylation data. (C) UMAP of 100 kb resolution CG DNA methylation profiles for the “Unknown” cluster identified in the 5 kb resolution clustering; cell-type labels are derived from the 100 kb resolution clustering. (D) UMAP of 10x Multiome RNA data using features selected for integration; cell-type labels are transferred from snm3C data. (E) Heatmap of –log₁₀(*P*) values from gene ontology (GO) enrichment analysis of genes upregulated in the “Unknown” cluster compared to other cell types. (F) Bar plot showing the fraction of each cell type in all samples (brain region + donor), grouped by brain region and disease state. (G) Bar plot of cell types whose proportions differ significantly among brain region and disease state groups.

We initially performed co-clustering with our previously published human brain snm3C-seq dataset^20^, using 100 kb binned mCG methylation profiles to transfer existing cell type labels to the current dataset (Extended Data Fig. 2A-B). To refine annotation accuracy, we conducted a second round of clustering using 5 kb genomic bins, calculating hypomethylation scores for each bin as outlined in the Methods. Clustering results at both 100 kb and 5 kb resolutions were largely concordant (Extended Data Fig. 2C), supporting the robustness of our annotations (Supplementary Table 2). After integrating data across donors, clustering based on 5 kb bins showed a uniform distribution across most metadata categories, except for a slight bias related to sequencing instruments (Extended Data Fig. 2D–L). We also identified a discordant cluster, labeled as “Unknown” (Figure 1B), consisting of cells inconsistently annotated between the two resolutions, suggesting potential heterogeneity or ambiguity within this population. To further investigate the identity of this group, we integrated snm3C-seq data with 10x multiome data by anchoring on cluster-enriched genes, aligning gene body methylation from snm3C-seq with gene expression from the multiome dataset. In the integrated analysis, the ‘Unknown’ cluster emerged as a distinct population (Figure 1D). Differential gene expression analysis revealed a marked upregulation of nervous system development–related functions in the “Unknown” cluster compared with all other cell types (Figure 1E), suggesting a progenitor-like cell state. We also observed substantial variation in cell type proportions across donors and brain regions, with oligodendrocytes (ODCs) and excitatory neurons showing the most pronounced changes (Figure 1F). To assess AD–control differences across regions, we performed an ANOVA, which revealed significant region-specific variation (Figure 1G).

### Global methylation difference and the identification of disease-associated cell state

We observed distinct global DNA methylation (mC) differences between AD and control samples across both neuronal and non-neuronal cell types, with region-specific patterns. In TC, one of the most vulnerable regions in AD, both neuronal and non-neuronal cells exhibited elevated global CG methylation levels in AD compared to controls (Figure 2A). In contrast, VC, a relatively spared region, showed reduced global methylation in neurons from AD samples, while non-neuronal cells displayed minimal differences (Figure 2A). To further investigate these trends at higher resolution, we quantified methylation levels genome-wide using 1 Mb bins across all samples. This analysis revealed that methylation differences were broadly distributed across the genome rather than being confined to specific loci (Figure 2B, Extended Data Fig. 3). Consistent with the global trends, TC samples showed widespread hypermethylation in both neuronal and glial populations. In VC, however, excitatory neurons exhibited genome-wide hypomethylation, while glial cells showed a mild but consistent hypermethylation in AD compared to controls (Figure 2B, Extended Data Fig. 3). These findings highlight the region– and cell type–specific nature of DNA methylation alterations in AD.

**Figure 2.**
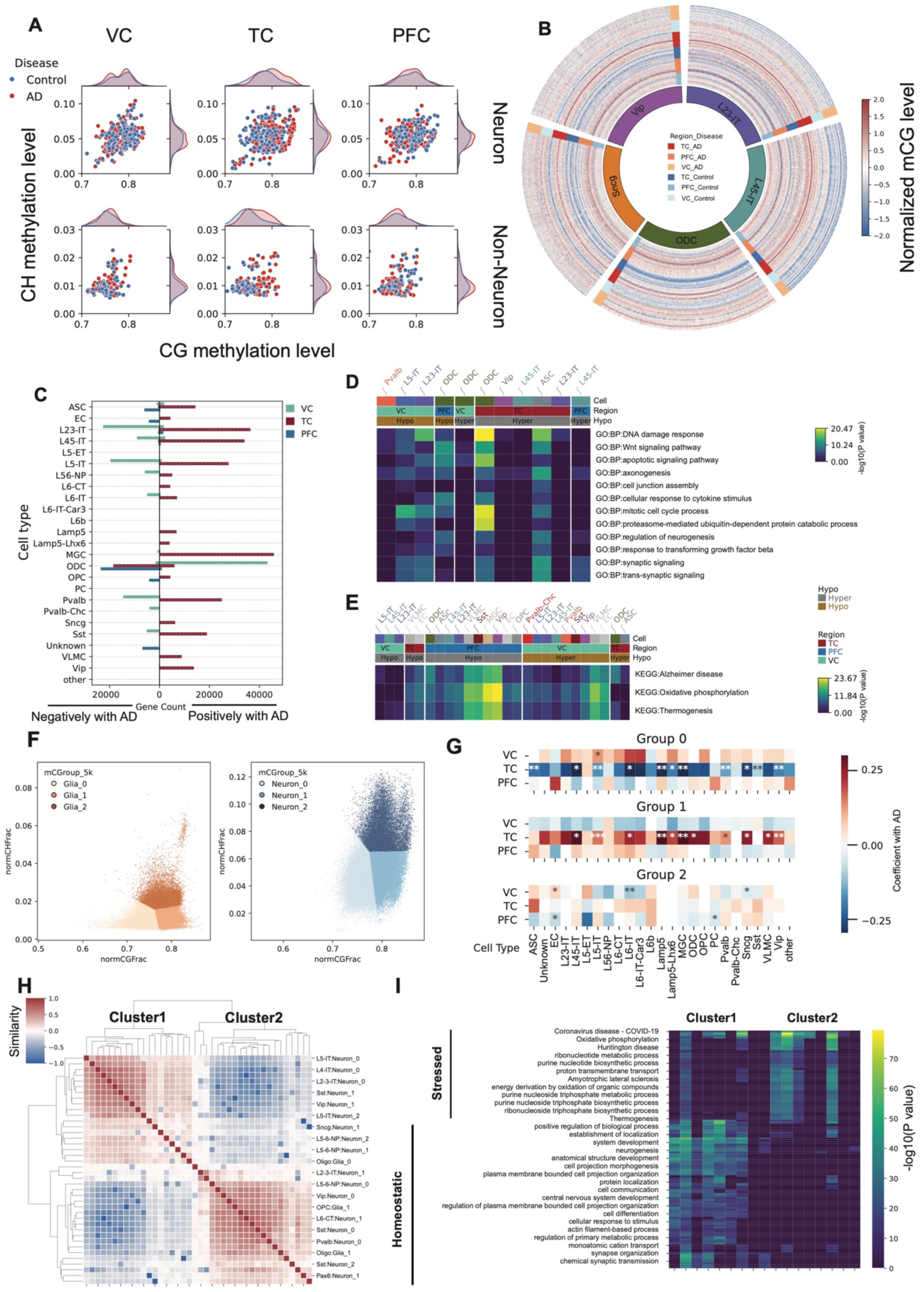
DNA methylation changes associated with Alzheimer’s disease (AD) across brain regions and cell types. (A) Scatter plot showing the median global CH and CG methylation levels for different brain region–disease state groups. (B) Circos plot of normalized genome-wide CG methylation levels in 1 Mb bins for five representative cell types (L23-IT, L45-IT, ODC, Sncg, Vip). (C) Bar plot showing the number of genes whose promoter CG methylation levels are positively (right) or negatively (left) associated with AD pathology. (D) Heatmap of –log₁₀(*P*) values from gene ontology (GO) enrichment analysis of genes associated with homeostatic functions. (E) Heatmap of –log₁₀(*P*) values from GO enrichment analysis of genes associated with AD-related stress responses. (F) Scatter plots showing single-cell global CH and CG methylation levels for non-neuronal cells (left) and neurons (right), colored by three subgroups defined by joint CH/CG methylation clustering. (G) Heatmaps showing Coefficients between subgroup-specific methylation features in each cell type and AD pathology (\**P* < 0.1,** P < 0.05, ***P < 0.01). (H) Clustered heatmap of similarities between subgroups across all cell types based on pairwise log₂ fold changes in differential expression analysis. (I) Heatmap of genes upregulated in either subgroup compared to other subgroups within each cell type, with color representing –log₁₀(*P*) values.

We identified genes whose methylation levels in each cell type were positively or negatively associated with Alzheimer’s disease (AD) in each brain region, using VC as a baseline in linear regression models (Methods). Consistent with the global hypermethylation observed in TC in AD, we found widespread gene hypermethylation across most cell types—except in oligodendrocytes (ODC), where VC already showed a highly methylated baseline (Figure 2B-C, Supplementary Table 3). Gene ontology enrichment revealed opposing patterns between VC and PFC/TC: genes that were hypermethylated in TC but hypomethylated in VC were enriched for homeostatic functions such as neurogenesis and axonogenesis, while genes that were hypomethylated in PFC and TC but hypermethylated in VC were enriched for stress-related functions, including Alzheimer’s disease and thermogenesis pathways (Figure 2D-E).

To distinguish whether global methylation differences across cell types reflect uniform shifts in methylation levels or changes in the proportions of subpopulations with distinct methylation states, we stratified both neurons and non-neurons into three groups based on CG and CH methylation: high CG/high CH (subgroup 2), high CG/low CH (subgroup 1), and low CG/low CH (subgroup 0). In TC, the proportions of these groups were significantly associated with AD status: the low-CG/low-CH group was negatively associated with AD, whereas the high-CG/low-CH group was positively associated. Notably, this pattern was reversed in VC, although most associations in this region were not statistically significant (Figure 2G). Subgroup 2 exhibited only weak correlations with AD across all three regions, suggesting that CH methylation in these cells may play a role in regulating their states. To further characterize the cell states associated with global methylation-defined subtypes, we transferred these group labels to the integrated 10x multiome dataset and identified differentially expressed genes (DEGs) between each pair of groups. We clustered the subtypes based on their pairwise log2 fold changes in gene expression, which revealed two major clusters of subtypes (Figure 2H). To further characterize these two clusters, we performed GO enrichment analysis on DEGs between subtypes within the same cell type across the two clusters. Genes upregulated in one cluster were consistently enriched for homeostatic functions, whereas those upregulated in the other were enriched for stress-related pathways (Figure 2I), including oxidative phosphorylation and neurodegenerative disease-associated processes. Based on these expression signatures, we defined the two clusters as representing *Homeostatic* and *Stressed* cell states, respectively. These states were observed across the majority of cell types and will be described in more detail in the following sections.

### 3D genome difference between AD and controls

We performed an additional clustering based on 100 kb contact matrices after imputation^21^, and transferred cell type labels from mC-defined cell types (Figure 3A). Interestingly, a substantial fraction of cells from the unknown cluster clustered with oligodendrocytes (ODCs) in this modality (Figure 3B), suggesting divergent cell identities between the DNAm and 3D genome modalities.

**Figure 3.**
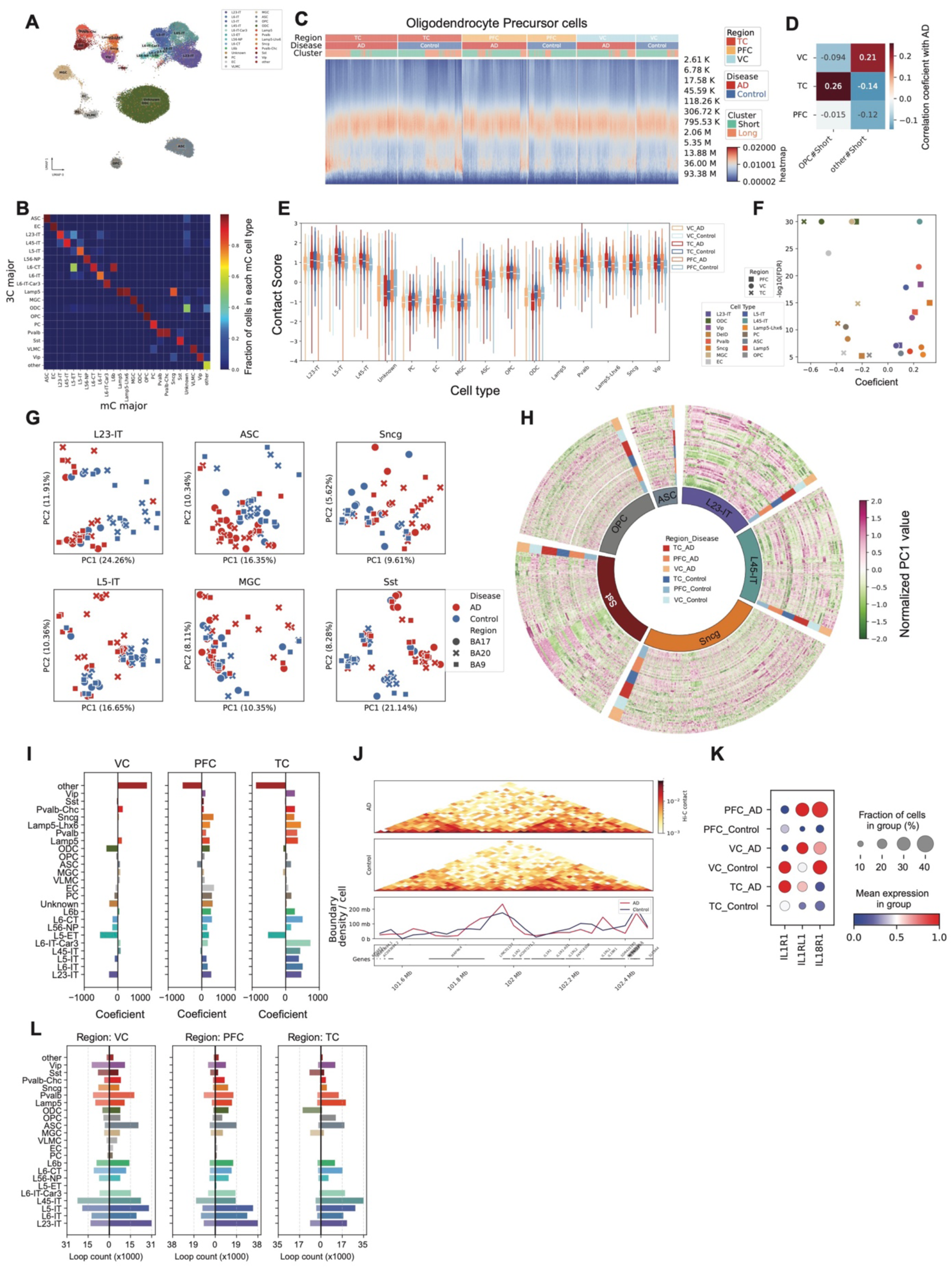
3D genome changes associated with Alzheimer’s disease (AD) across brain regions and cell types. (A) UMAP of single cells based on 100kb-resolution pairwise 3D genome contact maps. (B) Heatmap showing the proportion of DNA methylation–based cell type assignments within each 3D genome–derived cluster. (C) Heatmap showing the relationship between contact frequency and genomic distance in single cells. Colors indicate the fraction of contacts at each distance; annotations indicate brain region and disease state. Cells enriched in long-range contacts are classified as *Long*, and others as *Short*. (D) Violin plots of contact scores for each group in each cell type. (E) *P* values and coefficients showing associations between contact scores and AD pathology across groups and cell types. (F) Scatter plots of principal component analysis (PCA) on PC1 values from 3D genome compartment profiles. (G) Circos plot of normalized PC1 values across samples for genomic bins whose PC1 values are significantly associated (positively or negatively) with AD. (H) Bar plots of coefficients between the number of TAD boundaries and AD pathology in each brain region. (I) Heatmaps of normalized contact maps at the *IL1R1* and *IL18R1* loci in endothelial cells from the TC region in AD and control samples. Line plot shows boundary density in the same region at 100 kb resolution. (J) Dot plot of relative *IL1R1* and *IL18R1* expression levels in endothelial cells across brain regions and disease states. (K) Bar plots showing the number of differential chromatin loops between AD and control. Right, loops positively associated with AD; left, loops negatively associated with AD.

Contact probability versus distance curves revealed distinct chromatin folding patterns across brain regions and disease states. In TC, oligodendrocyte precursor cells (OPCs) from both AD and control samples showed a higher proportion of cells enriched in long-range contacts (Figure 3C). Ratios of short-or long-range contact varied in different cell types in AD and control in these three regions (Extended Data Fig. 4A). Interestingly, proportions of cells enriched in short-range contacts in OPC were associated with AD in TC, but the association is reversed in VC, suggesting region-specific alterations in chromatin architecture in AD. To quantify these patterns, we calculated a contact score for each cell, where lower scores indicate enrichment in long-range contacts and higher scores reflect enrichment in short-range contacts (Extended Data Fig. 4B). In non-neuronal cell types, AD cells exhibited lower contact scores compared to controls. In contrast, the opposite trend was observed in neurons (Figure 3D). Using a linear regression model to assess the relationship between contact score and AD status in each brain region, we found that AD was consistently associated with decreased contact scores in non-neurons and increased scores in neurons across all three regions (Figure 3E). This observation might indicate abnormal 3D genome states as neurons are enriched in short-range (higher contact score) contacts, while non-neurons are enriched in long-range (lower contact score) contacts (Extended Data Fig. 4A). To further relate this contact score to 3D genome organization within each cell type, we found it was positively correlated with the number of topologically associated domain (TAD) boundaries— particularly in non-neuronal and inhibitory neuronal populations (Extended Data Fig. 4C). Although the TAD analysis is described in detail later in the manuscript, we use those results here to establish this relationship.

We merged the contact matrices for each cell type and donor-region combination, using cell identities defined by DNA methylation profiles. To investigate compartment-level changes associated with AD across different cell types, we calculated compartment scores from the raw contact matrices for each aggregated sample. In several cell types, principal component analysis (PCA) of the compartment scores revealed clear separation between AD and control samples for some cell types (Figure 3F, Extended Data Fig. 5A). We further identified specific compartments whose activity was significantly associated with AD and observed widespread compartment shifts across multiple cell types (Figure 3G, Extended Data Fig. 5B, Supplementary Table 4), many of which were linked to corresponding gene expression changes.

To further interrogate chromatin architecture in AD, we identified TADs in individual cells using imputed contact matrices binned at 25 kb. The short-range contact score was positively correlated with the number of TAD boundaries within each cell type (Extended Data Fig. 4C), consistent with the idea that enriched short-range contacts accompany more sharply insulated TAD structures. Moreover, the correlation between per-cell boundary number and AD strengthened from VC to TC (Fig. 3H), indicating region-specific increases in chromatin insulation associated with AD. Although the total number of boundaries was positively associated with AD in TC and PFC, we also observed genomic intervals whose boundary density showed heterogeneous (both increased and decreased) associations with AD pathology (Supplementary Table 5).

To examine the genomic distribution of TAD boundaries, we calculated boundary densities using 1 Mb bins for each cell type and sample. Across both AD and control samples, PFC consistently exhibited the highest boundary density among the three brain regions. Additionally, we observed genome-wide differences in boundary density across several cell types, with particularly pronounced alterations in multiple neuronal populations in VC (Extended Data Fig. 6) and consistently lower boundary density in TC in both AD and control samples. Increased boundary density was associated with gene expression changes. For instance, at the *IL1RL1* and *IL18R1* locus, AD samples showed higher boundary density compared to controls in TC (Figure 3I), which coincided with elevated expression of these genes in endothelial cells (Figure 3J).

At the loop level, we observed widespread alterations associated with AD across all cell types and brain regions. In both PFC and TC, most cell types exhibited a greater number of gained loops in AD compared to controls. In contrast, VC showed a more balanced number of gained and lost loops in AD (Figure 3K, Supplementary Table 6), suggesting region-specific differences in 3D genome reorganization.

### Opposing epigenetic and transcriptomic associations in AD across brain regions

We identified AD-associated features using a linear regression model and found that the same feature, across three molecular modalities including DNA methylation, 3D genome and transcriptome, exhibited opposing associations with AD between VC and PFC/TC (Figure 4A). This opposing trend was most pronounced between VC and TC, with PFC displaying an intermediate pattern—consistent with the known gradient of regional vulnerability in AD. Interestingly, in most cell types except astrocytes (ASC), the genes showed hypomethylation and increased expression in AD in VC, with the opposite pattern observed in TC (Figure 4A, Supplementary Table 7).

**Figure 4.**
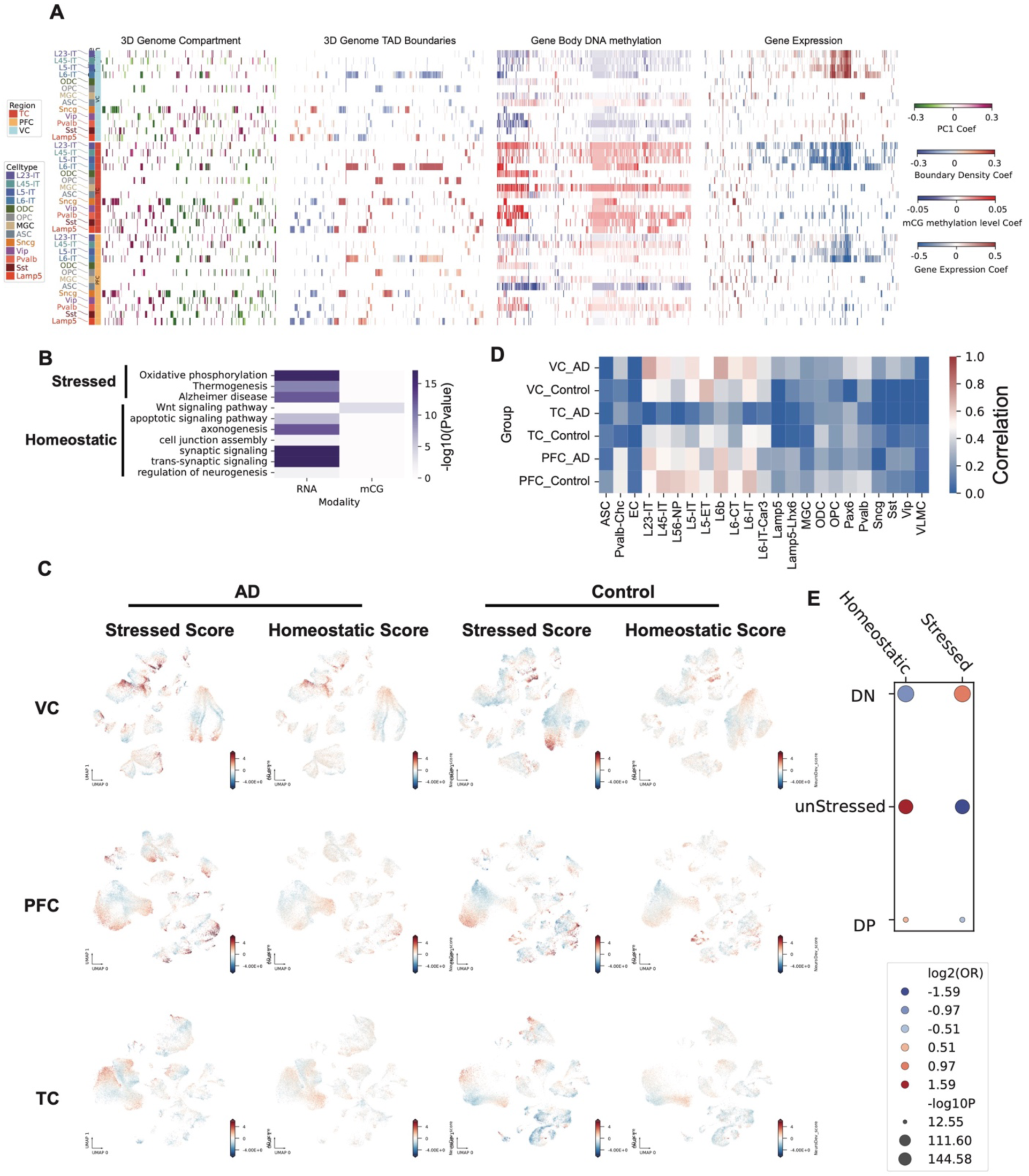
Opposing associations of multi-omic features with Alzheimer’s disease (AD) across brain regions. (A) Heatmap of coefficients for features from different modalities (DNA methylation, 3D genome, RNA) associated with AD pathology across brain regions and cell types. (B) Heatmap of Gene Ontology (GO) enrichment for genes similarly associated with AD in more than five cell types. Color indicates –log10(*P* value). (C) UMAP plots showing stressed scores (module score for genes in the GO term “Alzheimer’s disease”) and homeostatic scores (module score for genes in the GO term “neuron development”) for single cells from AD and control brains across three brain regions. (D) Heatmap of correlations between stressed scores and homeostatic scores across all cells in each cell type and brain region/disease state. (E) Scatter plot showing enrichment relationships between RNA-defined cell states and DNA methylation–defined states.

We further identified genes that were hypermethylated in TC and hypomethylated in VC in AD across multiple cell types. Similarly, we identified genes that consistently showed lower expression in TC and higher expression in VC in AD across cell types. These genes were enriched for homeostatic functions, including neuronal development and neurogenesis, as well as stress-associated functions such as Alzheimer’s disease (Figure 4B), mirroring the *Homeostatic* and *Stressed* cell states we defined based on DNA methylation.

We used scDRS^22^ to calculate a gene module score for each cell, using “Neurodevelopment” and “Alzheimer’s disease” genes as representative signatures of the Homeostatic and Stressed states, respectively. We observed substantial co-expression of the two modules in neurons, particularly in VC AD samples (Figure 4C). The Stressed score was high in PFC neurons in both control and AD, whereas both modules were low in TC, with control samples showing repression of the Stressed module (Figure 4C). The two module scores were highly correlated in excitatory neurons, except in AD samples from TC (Figure 4D). We next defined cell states based on the Stressed and Homeostatic scores and transferred these RNA-derived states to snm3C-seq cells through integration. Cells with both modules repressed (double negative, DN) were enriched in the Stressed cell state defined by DNA methylation, whereas cells with only the Stressed module repressed were enriched in the Homeostatic state defined earlier. Cells positive for both modules showed a slight enrichment in the Homeostatic state (Figure 4E).

### Putative deletions near telomeres in multiple cell types with the AD-associated state

To compare overall chromosomal contact patterns between cell states, we divided each arm of the autosomes into an equal number of bins and calculated the average contact frequency in both stressed and homeostatic states. The log₂ fold change in contact revealed a consistent reduction in telomeric contacts in the AD-associated state across multiple cell types (Figure 5A, Extended Data Fig. 7A). To investigate this effect further, we visualized contact maps at the telomeric end of chromosome 8 in L23-IT cells and observed widespread contact loss in the stressed state across all three brain regions (Figure 5B), suggesting structural alterations at chromosome ends under stress. To comprehensively assess contact loss across the genome, we calculated total normalized contacts in each 100 kb bin and filtered out bins with no contacts across all cell types and states. We found that numerous bins lacked contacts specifically in the AD-associated state across cell types and regions (Figure 5C, Supplementary Table 8). These putative deletion regions were predominantly located near telomeres, as shown by their genomic distribution (Figure 5D). We identified genes located within these deleted regions. We found that they were significantly enriched for functions related to response to stimulus and signaling pathways (Figure 5E), suggesting that telomere-proximal chromatin alterations in the AD-associated state may impact stress-responsive gene regulation. Notably, the number of bins lacking contacts was significantly higher in the AD-associated cell states (Figure 5F, Extended Data Fig. 7B), which might be related to the association between shorter telomeres and AD pathology^23–26^.

**Figure 5.**
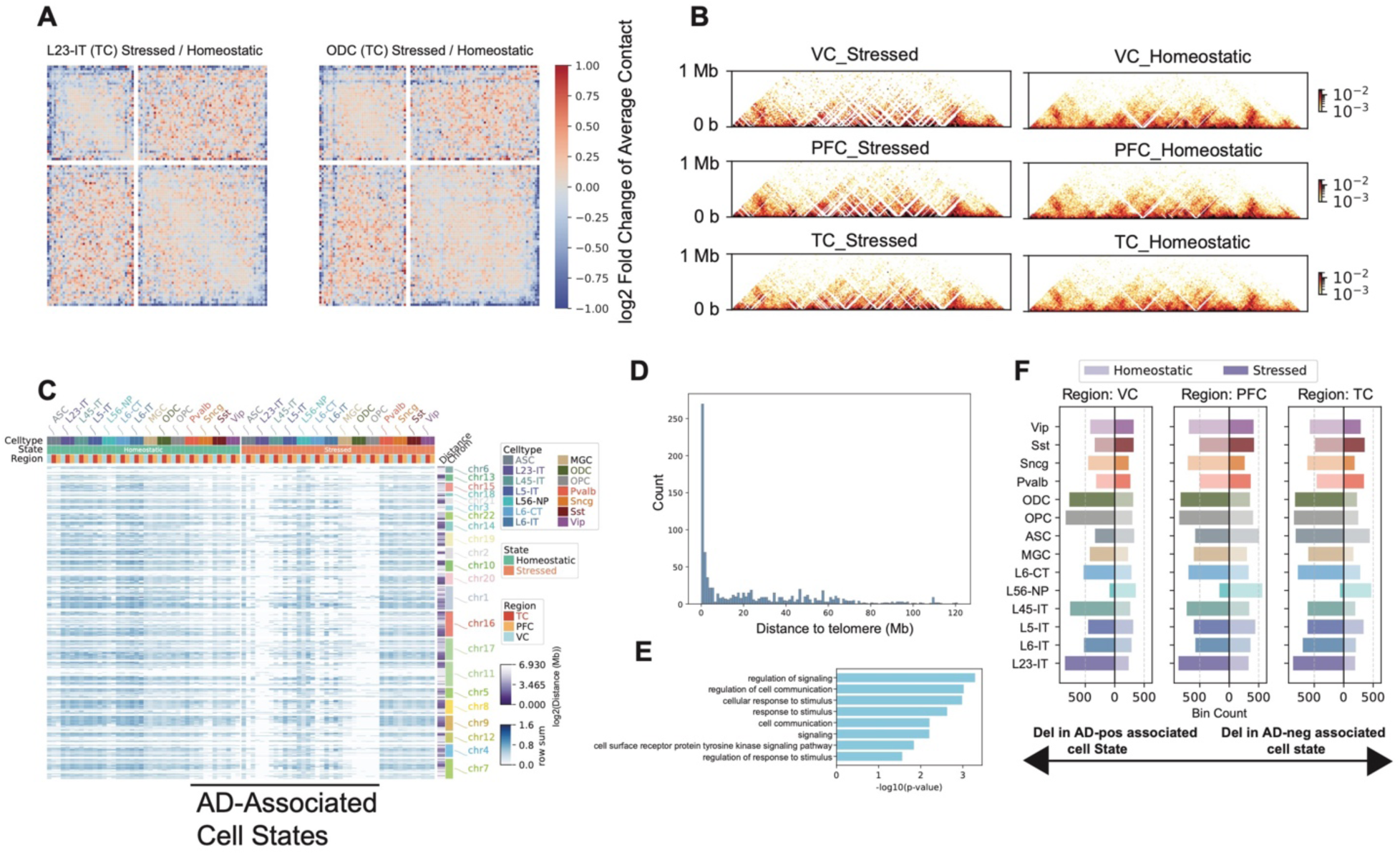
Putative telomere deletions in stressed states across cell types. (A) Heatmap showing the division in averaged obs/exp contact maps across all chromosomes with scaled p– and q-arms in TC for L23-IT and ODC cell types. (B) Heatmaps of normalized contact frequencies in a 5 Mb region at the end of chromosome 8, separated by stressed and homeostatic states across three brain regions. (C) Heatmap of row sums of normalized contacts for bins defined as putative deletions. Column annotations indicate cell type, cell state (stressed or homeostatic), and brain region. Row annotations indicate chromosome identity and the genomic distance of the deletion to the nearest telomere. (D) Histogram showing the number of putative deletions (100 kb bins) at different distances from telomeres. (E) Bar plot of GO enrichment for genes located within putative deletions. (F) Bar plots showing the number of putative deletions in stressed (left) and homeostatic (right) states for each cell type.

### Epigenomic signatures of Stressed and Homeostatic states

To characterize the epigenomic features of homeostatic and stressed states identified in each cell type, we performed comprehensive comparisons spanning 3D genome organization, DNA methylation, and gene expression. We first compared the compartmentalization patterns of the two states within each cell type by generating saddle plots, which quantify the average interactions between bins with similar PC1 values. Subtracting the homeostatic matrix from the stressed matrix revealed that, in most cell types across the three regions, AA interactions were strengthened under stress. The only exception was the synuclein gamma–positive neuron (Sncg) population, which did not show this increase in PFC and TC (Figure 6A).

**Figure 6.**
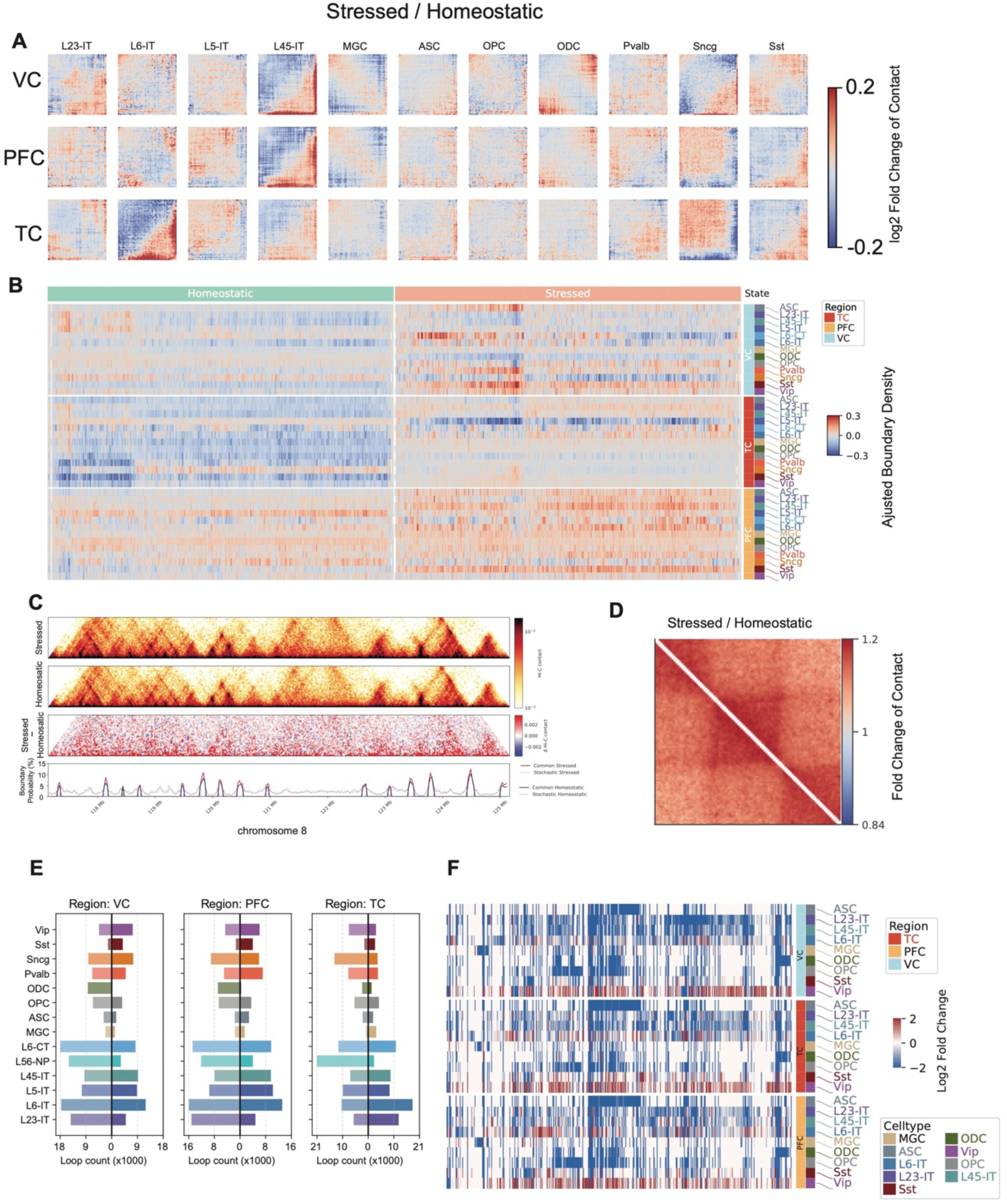
Epigenomic features of Stressed and Homeostatic states across cell types. (A) Heatmaps show the division (*Stressed / Homeostatic*) of compartment saddle plot matrices. (B) Heatmap shows the adjusted boundary densities in stressed and homeostatic states across brain regions and cell types. (C) Genome browser view shows the contact heatmaps of stressed, homeostatic, and the subtraction (*Stressed – Homeostatic*) between them, as well as the stochastic and common boundaries in these two states in an 8 Mb region on chromosome 8. (D) Division (*Stressed / Homeostatic*) of pileup matrices centered on TADs between stressed and homeostatic states in L23-IT in the TC region. (E) Barplots show the number of differential loops between stressed and homeostatic states in all cell types and three brain regions. Right side of x = 0 are the number of loops gained in stressed, while the left side are gained in homeostatic. (F) Heatmaps show the log₂ fold changes (*Stressed / Homeostatic*) across cell types and brain regions.

TAD organization also exhibits global differences between the two states. In each brain region and across multiple cell types, we observed a consistently higher genome-wide TAD boundary density in the stressed state compared to the homeostatic state (Figure 6B), indicating widespread reorganization of 3D genome architecture under stress. To determine whether stochastic or shared boundaries drove this increase, we quantified both. We found that the densities of both stochastic and common boundaries were elevated in the stressed state (Figure 6C) in Layer 2/3 Intratelencephalic Neurons (L23-IT) in TC, accompanied by an increase in short-range chromatin contacts. To further assess whether this increase in short-range contacts reflects local or genome-wide changes, we performed TAD-centered pileup analysis by rescaling TADs to a uniform size with 100 kb flanking regions on both sides. This analysis revealed that contact frequencies were elevated in the stressed state both *within* TADs and *across* TAD boundaries (Figure 6D) in L23-IT, supporting a global shift toward increased local chromatin insulation and compaction under stress. Pileup analysis centered on TADs of other cell types and regions shows that short-range contacts are either increased or decreased between the two cell states (Extended Data Fig. 8).

We also observed widespread changes in chromatin loops between the stressed and homeostatic states, with the most prominent alterations occurring in excitatory neurons (Figure 6E, Supplementary Table 9). Differential gene expression analysis between Homeostatic and Stressed cell states showed that genes are globally repressed in excitatory neurons and non-neurons (Figure 6F).

## Discussion

Epigenomic dysregulation in AD was revealed about two decades ago^15,27,28^, and epigenetic drugs have been tested in models of AD^17,29–31^. More recently, advances in single-cell chromatin accessibility profiling have introduced the concept of “epigenome erosion**”**, describing a progressive loss of chromatin accessibility patterns associated with cell identity^10,11^. We performed single-nucleus DNA methylation and chromatin conformation profiling across multiple brain regions—the visual cortex (VC), prefrontal cortex (PFC), and temporal cortex (TC)—providing a multimodal view that complements traditional single-modality studies. Our study revealed global changes in both DNA methylation and the 3D genome in AD with changes in VC and TC opposing each other. More importantly, we defined *Homeostatic* and *Stressed* states for multiple cell types, which showed consistent changes in all modalities. Intriguingly, we found there are more deletions near telomeres in AD-associated states across multiple cell types.

Consistent with the loss of chromatin accessibility in AD^10,11^, we observed widespread hypermethylation in the prefrontal cortex (PFC) and temporal cortex (TC) across most cell types. In contrast, the visual cortex (VC) showed the opposite trend, with global hypomethylation in AD, underscoring the region-specific nature of epigenomic dysregulation. Similarly, the number of chromatin boundaries associated with AD increased in PFC and TC but decreased in VC. These findings suggest that regions relatively spared in AD should not be overlooked: their apparent resilience may reflect not an absence of molecular alterations, but the presence of protective epigenetic changes that help preserve function. Although the associations in VC run counter to those in PFC and TC, whether these changes are protective remains to be determined. Such features provide important clues for prevention strategies aimed at more vulnerable brain regions.

In chromatin accessibility-based studies, cell identity loss^10^ and widespread epigenomic relaxation and information loss^11^ were linked to cognitive resilience. Based on our genome-wide analysis, we identified two distinct cell states—Homeostatic and Stressed—across most cell types. The AD-associated state consistently exhibits increased putative genomic deletions near telomeres, a higher density of TAD boundaries, and broad repression of gene expression, particularly in excitatory neurons and non-neuronal cells. Multiple studies have reported an association between shorter telomere length and AD^23–26^, although a causal relationship has yet to be established. This phenomenon may be linked to deletions occurring near telomeres in disease-associated cellular states, potentially driven by stress-induced telomere shortening^32^ given their critical role in protecting chromosome ends. Recent studies have uncovered a non-canonical role for DNMT3A in regulating telomerase activity and preserving genome integrity^33^, suggesting a possible mechanistic link between the global methylation changes we observe and telomere deletions. However, in our data, DNMT3A expression does not correlate with the number of deletions, indicating that additional factors are likely involved. Evidence of epigenetic alterations, particularly a global increase in DNA methylation, has previously been reported in the context of telomere shortening^34,35^.

The “Unknown” cluster, enriched for genes involved in nervous system development, is of particular interest. Although these cells are annotated as different cell types based on 100 kb methylation patterns, they appear similar at 5 kb resolution, suggesting a potential cell-type transition linked to neurogenesis. This notion is further supported by the observed discordance in cell identity between DNA methylation and 3D genome features. While a detailed investigation lies beyond the scope of the present study, this intriguing observation warrants future exploration.

### Limitations of the study

While our results provide a cell-type– and region-specific multi-omic view of AD brains, the limited sample size (11 AD and 9 control), restricted brain regions examined (VC, PFC, and TC), and narrow genetic background of donors mean we are still far from achieving a comprehensive, multi-modal characterization of AD brains. Owing to the scope of this study, we did not investigate the underlying mechanisms of the features observed in AD brains; future mechanistic studies will be necessary to elucidate these processes. Although we detected deletions in regions adjacent to telomeres, we were unable to directly assess telomere length in these cell states. Future studies will be needed to determine whether these deletions are driven by telomere shortening under stress.

## Methods

### Data Generation

#### Human tissue collection and neuropathological evaluation

Post-mortem human brain tissues were obtained from the Massachusetts Alzheimer’s Disease Research Center (MADRC, P30AG062421) with informed consent from patients or their next of kin, and according to protocols approved by the Partners Human Research Committee. Each donated brain was bisected into two hemispheres at autopsy. One hemisphere was immersion-fixed in 10% formalin, paraffin-embedded, and sectioned for neuropathologic evaluation. The contralateral hemisphere was coronally sectioned, flash frozen on dry ice, and stored at –80 °C. For this study, frozen cortical tissues were dissected from the left hemisphere and processed for downstream molecular analyses.

Neuropathologic assessment was performed by a board-certified neuropathologist at MADRC in accordance with the National Institute on Aging–Alzheimer’s Association (NIA-AA) guidelines^36,37^. Evaluation included immunohistochemistry for Aβ, tau, α-synuclein, and phosphorylated TDP-43, as well as assessment for vascular pathology. Clinical and demographic information (age at death, sex, postmortem interval, and APOE genotype) was collected where available and is summarized in Supplementary Table 1.

#### Sample transfer

Brain tissue samples were transported on dry ice from Massachusetts General Hospital (MGH) to the Salk Institute and promptly stored at –80 °C until further processing.

#### Nuclei isolation and Fluorescence-Activated Nuclei Sorting (FANS)

Nuclei isolation was performed following a standard protocol (dx.doi.org/10.17504/protocols.io.y6rfzd6). Briefly, single nuclei were stained with Alexa Fluor 488–conjugated anti-NeuN antibody (A60, monoclonal, MAB377X, Millipore; 1:500 dilution) and Hoechst 33342 (62249, Thermo Fisher), then subjected to fluorescence-activated nuclei sorting (FANS) on a BD Influx sorter in single-cell (one-drop single) mode. For each 384-well plate, NeuN⁺ (488⁺) nuclei were sorted into columns 6–24, and NeuN⁻ (488⁻) nuclei into columns 1–5, yielding an approximate 3.8:1 ratio of NeuN⁺ to NeuN⁻ nuclei. The snm3C-seq workflow incorporated in situ 3C treatment during nuclei preparation, enabling simultaneous capture of chromatin conformation. These steps were carried out using the Arima-3C BETA kit (Arima Genomics).

#### snm3C-seq library preparation and sequencing

Snm3C-seq samples followed the library preparation protocol described previouslydescribed^19,38^(details:https://www.protocols.io/view/snm3c-seq3-kqdg3x6ezg25/v1). This protocol was automated using a Beckman Biomek i7 and Tecan Freedom Evo instrument to facilitate large-scale applications. The snm3C-seq libraries were sequenced on an Illumina NovaSeq 6000 and NovaSeq X instruments in two batches, using one S4 flow cell per 16 384-well plates and using 150 bp paired-end mode. The following software were used during this process: BD Influx (v.1.2.0.142; for flow cytometry), Freedom EVOware (v.2.7; for library preparation), Illumina MiSeq control (v.3.1.0.13) and NovaSeq 6000 and X control (v.1.6.0/RTA, v.3.4.4; for sequencing), and Olympus cellSens Dimension 1.8 (for image acquisition).

## Data Analysis

### Mapping and Quality Control

Mapping of snm3C-seq data was performed using the YAP pipeline (cemba-data package, v1.6.8) as previously described^38^. The main workflow included: (1) demultiplexing FASTQ files into single cells (cutadapt^39^, v2.10); (2) read-level quality control (QC); (3) mapping for snm3C-seq (bismark^40^, v0.20; bowtie1^41^, v1.3.1); (4) BAM file processing and QC (samtools^42^, v1.9; Picard, v3.0.0); (5) methylome profile generation (allcools, v1.0.23); and (6) chromatin contact calling. Detailed Snakemake^43^ pipeline files are provided in the *Code availability* section. All reads were aligned to the combined reference of human Genome Reference Consortium build 38 (GRCh38) and lambda genome, with gene and transcript annotations based on a modified GENCODE v35 GTF file.

Primary QC for DNA methylome data applied the following thresholds: (1) overall mCCC level < 0.05; (2) overall mCH level < 0.2; (3) overall mCG level > 0.5; (4) total final reads between 0.5 M and 10 M; and (5) bismark mapping rate > 0.5. The mCCC level was used to estimate the upper bound of the bisulfite non-conversion rate at the cell level. Lambda DNA spike-in methylation was also quantified to estimate the non-conversion rate per sample, with all samples showing values < 0.01 (Extended Data Fig. 2i). Relatively loose cutoffs were chosen at this stage to avoid loss of valid cells or clusters, with further QC performed during clustering to identify potential doublets and low-quality cells.

For the 3C modality in snm3C-seq cells, an additional requirement was applied: > 50,000 cis long-range contacts (anchors > 2,500 bp apart).

### Methylome clustering analysis

After mapping, single-cell DNA methylome profiles from the snm3C-seq datasets were stored in the “all cytosine” (ALLC) format—a tab-separated table compressed and indexed using bgzip/tabix. The *generate-dataset* command in the ALLCools package was used to generate a methylome cell-by-feature tensor dataset (MCDS). For clustering analysis, we used non-overlapping 100-kb genomic bins (chrom100k) from the hg38 genome. Gene body regions were used for clustering annotation and for integration with the companion 10x multiome dataset. Non-overlapping 5-kb genomic bins (chrom5k) were used for integration with the chromatin accessibility dataset. Details of the integration analyses are provided below.

### Co-clustering with Human Brain Atlas data

We used 100-kb (chrom100k) CG and CH DNA methylation profiles of the cells that passed DNA methylation QC to co-cluster our AD snm3C-seq data with previously published human brain snm3C-seq datasets^20^. We did some filtering and selection on the 100 kb bins: (1) bins should have > 500 and < 3000 coverage, (2) bins have less than 20% overlap with hg38 blacklist v2 regions, (3) bins should be on autosomes, ‘chrX’, ‘chrY’ and ‘chrM’ bins are removed, (4) top 20000 highly variable features were selected. Dimensionality reduction was performed jointly on the two datasets for CG and CH profiles separately. For CG and CH, the top 63 and 95 principal components (PCs), respectively—determined by a difference significance threshold of 0.1 relative to the last PC—were concatenated for clustering. Nearest-neighbor detection and Leiden^44^ clustering were performed in Scanpy^45^ with *n_neighbors = 25* and *resolution = 1.5*; all other parameters were left at their default settings. Batch effects between the reference human brain atlas and AD snm3C-seq data were corrected using Harmony^46^. Cell type labels were assigned to Leiden clusters based on reference dataset annotations and used to annotate our AD snm3C-seq cells.

We also performed an independent clustering using only CG methylation profiles from the AD snm3C-seq data, applying the same parameters. Both CG– and CH-based clustering produced comparable results. Final cell type assignments were determined using a majority-vote approach, in which each Leiden cluster was assigned to the cell type represented by the largest number of cells within that cluster.

### Methylome clustering using 5 kb bins

We did another higher resolution clustering using 5 kb bins, applying the same feature filtering criteria as described above. We first quantified the CG methylation level of the 5 kb bins across whole genome, then calculated the hypomethylation score of each bin within each cell according to the method below:

Hypo-score measures the likelihood of observing greater than m methylated reads under the assumption that methylation follows the binomial distribution with parameters *c* and *p*.

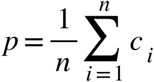

m is the observed number of methylated count for region i, c is the coverage (total count) covering region i and n is the total number of 5kb bin regions, p is the expected probability of methylation for this cell.

Let’s assume

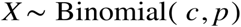

then for each 5 kb bin,

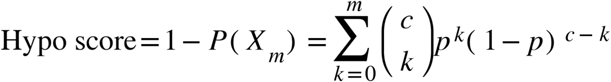

The calculation of hypo score was implemented in ALLCools (https://lhqing.github.io/ALLCools/intro.html) using scipy^47^

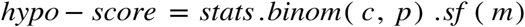

The hypomethylation score matrix was binarized using a cutoff of 0.9, such that bins with scores > 0.9 were assigned a value of 1 and all others a value of 0. Clustering was performed using the same parameters as in the previous analysis. When using 5-kb bins, clustering revealed high donor heterogeneity; Harmony was therefore applied to integrate data across donors. Cell type labels were assigned by majority vote, based on the cell identities determined from the 100-kb bin clustering.

### Integration with 10x multiome data

Based on the high-efficiency framework developed in previous study^38^, based on the Seurat R package Canonical Correlation Analysis (CCA)^48,49^ integration algorithm, to perform atlas-scale data integration with millions of cells. The framework consisted of three main steps to align two datasets in a shared space: (1) applying dimensionality reduction to generate embeddings for both datasets in the same space; (2) CCA to capture shared variance across cells between datasets and identify anchors, defined as five mutual nearest neighbours between the two datasets; and (3) aligning the low-dimensional representations of the two datasets using these anchors. Genes were used as features to integrate methylome and transcriptome profiles.

To integrate our snm3C-seq dataset with 10x multiome, 10x cell gene expression values were normalized by dividing each cell’s unique molecular identifier (UMI) count by the total UMI count for that cell, multiplying by the mean total UMI count across all cells, and log-transforming the result. For snm3C-seq cells, posterior gene body mC levels were used. Cluster-enriched genes (CEGs) were identified for each cell subclass and cluster using snm3C-seq methylation data, and only CEGs with mC variance > 0.05. Because gene body methylation is negatively correlated with gene expression^38^, mC levels were sign-reversed prior to integration.

A principal component analysis (PCA) model was fit to the snm3C-seq cells, and the 10x cells were projected into the same space. PCs were normalized by dividing by the singular value of each dimension to prevent dominance by the first few PCs.

For anchor identification, the CEG matrices of snm3C-seq and scRNA-seq cells were Z-score scaled across cells, yielding matrices XXX (snm3C-seq: cell × CEG) and YYY (10x: cell × CEG). Canonical correlation analysis (CCA) was applied to find a shared low-dimensional embedding, solved by singular value decomposition of XtYX^\top YXtY. The resulting canonical variates UUU (snm3C-seq) and VVV (scRNA-seq) were row-normalized (L2 norm) and used to identify five mutual nearest neighbours as anchors, scored following the Seurat^49^ method.

Integration was then performed in PC space, with the PCs of the 10x dataset (query) projected onto the PCs of the snm3C-seq dataset (reference), while keeping the reference PCs unchanged. For each RNA–snm3C-seq anchor pair, the vector difference between their PCs was treated as a bias. For each 10x query cell, a weighted average bias vector was computed from its 100 nearest anchors in RNA PC space, with closer anchors receiving higher weights. This bias was applied to align 10x cells into the snm3C-seq space. And the cell type labels from snmc3C-seq data were transferred to the 10x multiome cells.

### Cell proportions across donors and regions

The proportions of cell types for each donor and brain region were calculated (Fig. 1f), and differences in distribution were tested using ANOVA. Significantly different cell types were then plotted (Fig. 1g).

### Cell-type-level DNA methylation analysis associated with AD

#### Whole-genome DNA methylation levels

We grouped the cells by the cell type labels and donor/brain region, and calculated the median CH and CG DNA methylation levels. The CH and CG DNA methylation levels of neuron and non-neuron cell types were plotted separately.

We merged the cells to pseudo-bulk by cell type/donor/brain region, and calculated the average DNA methylation levels across the whole genome in 1 Mb bins. Since we did sequencing using two different sequencers, we noticed a batch effect between the two instruments by sequencing the same library (Extended Data Fig. 3A-B). To remove variation attributable to technical batch while retaining biological covariate effects, we applied per-feature ordinary least squares (OLS) regression. For each genomic bin, the bin-level values were regressed on batch, age, sex, brain region, disease status, and donor identity, with categorical variables one-hot encoded. The fitted contribution of the batch terms was subtracted from the observed values, leaving residuals adjusted for batch but preserving variation explained by the other covariates. This procedure was applied independently to each bin, producing a corrected matrix for downstream analyses. We visualized the corrected DNA methylation levels by normalizing the CG methylation across samples, and plotted with pycirclize (https://github.com/moshi4/pyCirclize).

#### Differentially methylated genes analysis

We corrected batch effects at the gene level using per-gene ordinary least squares (OLS) regression on the cell metadata. For each gene, the vector of per-cell CG methylation values *yg*\*mathbf*{*y*}_*gyg* (rows = cells, columns = genes in the AnnData object) was modeled as

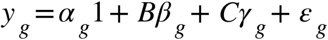

where Bcontains one-hot encoded batch indicators and C contains one-hot/continuous covariates (Age, Sex, Region, Disease, Donor). After fitting via OLS, we computed the fitted batch component 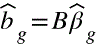 and subtracted it from the observed values to obtain batch-adjusted methylation:

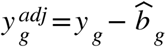

This procedure removes variation attributable to batch while preserving effects associated with the specified biological covariates and the intercept. Categorical variables were one-hot encoded with a reference level (drop-first), so the adjustment aligns all cells to the reference batch baseline.

##### Implementation details

The design matrix was constructed once from adata.obs and reused for all genes. OLS was fit independently per gene, iterating over columns of adata.X. For sparse inputs, the matrix was densified prior to fitting. The resulting batch-corrected matrix was written to a new AnnData object with original obs/var preserved for downstream analysis.

We tested per-gene associations between CG methylation and clinical/biological covariates using linear models fit separately within each cell type. For each cell type, we read the gene-by-cell AnnData matrix (batch-corrected upstream) and, for each gene *g*, modeled per-cell methylation values as:

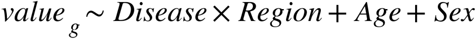

using ordinary least squares (statsmodels). Categorical variables were encoded with explicit reference levels: Disease (Control reference, AD as contrast), Region (VC reference; PFC and TC contrasts), and Sex (Female reference; Male contrast). The interaction term (^Disease × Region^) tests region-specific disease effects.

Quality filters excluded genes with fewer than 5 unique donors represented or with entirely missing values. For each gene, we extracted coefficient estimates and P-values for the AD main effect, region contrasts (PFC, TC), Sex, Age, and Disease×Region interactions (AD×PFC, AD×TC). Multiple testing correction was performed separately for each term across genes using the Benjamini–Hochberg false discovery rate (FDR). The resulting table reports, per gene, effect sizes, raw P-values, and FDR-adjusted Q-values for all tested terms. Parallel processing (Python multiprocessing) was used to iterate over genes, and outputs were written per 10,000-gene chunk. FDRs were calculated by merging the results for each cell type. Genes deemed “DMG” in downstream analyses were selected based on P values (<0.05) and absolute coefficient (>0.01). Gene Ontology (GO) enrichment of the DMGs were performed with GProfiler^50^, and a heatmap plotted with PyComplexHeatmap^51^.

#### Definition of Homeostatic and Stressed Cell States

The global methylation levels for both CG and CH contexts, measured from the same library on two Illumina sequencing instruments, exhibited a strong linear relationship (Extended Data Fig. 3A–B). We used this relationship to normalize the global methylation levels of all cells across instruments, thereby correcting instrument-specific batch effects. Cells were first divided into neuronal and non-neuronal groups, and CG and CH methylation levels were normalized separately within each group. We then applied *k*-means clustering to the normalized values, partitioning the cells into three clusters, corresponding to low CH, low CH (group 0), high CG, low CH (group 1) and high CG, high CH (group 2).

We tested the associations of these three groups defined by global DNA methylation level with AD pathology, using a similar linear regression model we used in DMG identification. We modeled per-feature cell ratios using ordinary least squares (OLS) with diagnosis, region, age, and sex as predictors, including a diagnosis-by-region interaction. Analyses were performed either across all cells or within a specified cell type. For each feature fff, rows corresponding to that feature were subset and the following model was fit:

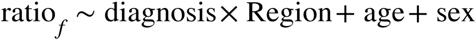

with categorical encodings: diagnosis (Control reference; AD contrast), Region (VC reference; PFC and TC contrasts), and sex (Female reference; Male contrast). Features represented by fewer than 5 unique donors were excluded. From each model we extracted effect sizes and P-values for the AD main effect, region contrasts (PFC, TC), age, sex, and the AD×Region interaction terms (AD×PFC, AD×TC). Multiple testing corrections used the Benjamini–Hochberg procedure applied per term across features to obtain FDR-adjusted Q-values. Computations were parallelized over features using Python’s multiprocessing.

##### Interpretation

The AD main-effect coefficient tests the difference in mean ratio between AD and Control in the reference region (VC), adjusting for age and sex. Interaction terms test whether the AD–Control difference differs in PFC or TC relative to VC.

We transferred the group labels (0, 1, 2) for each cell type to the 10x multiome cells using the integration results described above. Pairwise differential expressed genes (DEGs) between each two of the groups were identified with Seurat^52^ using MAST^53^, with “nFeature_RNA”, “Age”, “Sex”, “Region”, “Disease” as covariates. Pairwise log2 fold changes (log2FC) were used to cluster the groups defined with global methylation from all cell types. We cluster the subtypes (groups in each cell type) using the following method:

##### Input

A matrix of genes × comparisons (log2FC matrix), where each column encodes a within–cell-type contrast as celltype:SubtypeA_vs_SubtypeB.

### Procedure

1. Subtype set. We parsed all column names to enumerate the unique subtypes for each cell type (e.g., celltype:SubtypeA, celltype:SubtypeB).
2. Gene-wise directed adjacency matrices. For each gene ggg, we built a square matrix **A**^(g)^ indexed by subtypes. For every comparison*^S^_i_* vs *^S^_j_* with log2FC*^f^_g_*.
3. Tensor assembly and dimensionality reduction. We stacked {**A**^(g)^} over genes into a 3-D tensor [subtype×subtype×gene] ^subtype×subtype×gene^, flattened to 2-D, and applied PCA (50 components). The PCA scores were reshaped back to [subtype×subtype×50].
4. Subtype embeddings. For each subtype SSS, we summarized its PCA-transformed outgoing contrast profile by averaging across its row:

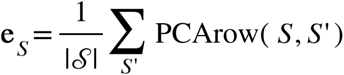

yielding a 50-dimensional embedding per subtype that reflects its gene-wise differential pattern against all peers.
5. **Similarity and clustering.** We computed cosine similarity between subtype embeddings to obtain **K** with:

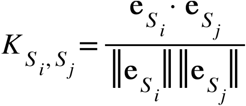

converted it to a distance matrix 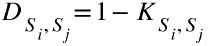, and performed hierarchical clustering (average linkage). We visualized the clustered similarity heatmap with the corresponding dendrograms.

The subtypes were grouped into two major clusters based on their similarity profiles. For each cluster, we identified genes uniquely upregulated within that cluster by intersecting cluster membership with differentially expressed genes (DEGs) from pairwise subtype comparisons within each cell type. Gene Ontology (GO) enrichment analysis was performed on these cluster-specific genes. Based on the GO terms consistently enriched across subtypes within each cluster, we annotated the two clusters as representing **Homeostatic** and **Stressed** states.

### Cell type level 3D genome analysis associated with AD

The chromatin contact data were generated in the YAP mapping pipeline as described above. We filtered out the contacts that overlapped with hg38 v2 blacklist, and another 2D blacklist^20^ which are mostly mis-aligned multimapping reads not filtered out from bismark bowtie1 mapping. Starting with these filtered contacts, we analyzed the 3D genome data in following procedures:

#### 1. Contact imputation in single cells

We imputed the contacts at three different resolutions for each single cell: 10 kb, 25 kb, and 100 kb using scHiCluster^21^ with the following parameters: ––batch_size 1536 ––pad 1 ––cpu_per_job 30 ––chr1 1 ––pos1 2 ––chr2 5 ––pos2 6. For 100 kb bin imputation, we use “––output_dist 500000000 ––window_size 500000000 ––step_size 500000000”, for 25 kb bin imputation we used “––output_dist 5050000 ––window_size 500000000 –– step_size 500000000”, and for 10 kb imputation, we used “––output_dist 5050000 –– window_size 30000000 ––step_size 10000000”.

#### 2. 3D genome embedding with 100 kb contacts

Three-dimensional (3D) genome embeddings were generated using the embedding function of scHiCluster applied to the imputed 100 kb contact matrices. Embeddings from all autosomes (excluding chromosomes X, Y, and M) were concatenated and stored in an AnnData object. Clustering of the combined embeddings was performed using the Leiden algorithm with parameters n_neighbors = 20 and resolution = 2. Cell-type labels for the resulting 3D genome clusters were assigned by majority voting based on DNA methylation–derived annotations. A confusion matrix was generated to compare DNA methylation–based and 3D genome–based clustering, calculated as the fraction of DNA methylation–defined cells assigned to each 3D genome cell type.

#### 3. Chromatin contacts versus genomic distance

The relationship between number of contacts and genomic distance was calculated with the “contact-distance” function in scHiCluster for each single cell. The contact density at different genomic distance heatmaps were plotted with PyComplexHeatmap. A Gaussian kernel was constructed over the genomic distance axis, centered at the midpoint of the distance range and with a standard deviation of one-sixth the total number of bins. The kernel was normalized to sum to 1. For each cell *c*, an enrichment score was computed as the weighted average of the contact–distance ratios across bins, using the Gaussian kernel as weights:

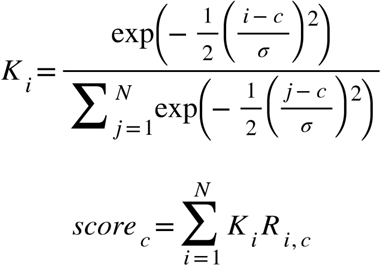

where *R*_*ic*_, is the ratio at bin *i* for cell *c*, and *K*_*i*_ is the corresponding kernel weight. Scores were optionally standardized (z-scored) across cells. Principal component analysis (PCA) was applied to reduce the data to two components, and *k*-means clustering (k = 2) was performed in this PCA space to group cells with similar decay profiles. The correlation of the score and “short” or “long” range contact enriched cells with AD pathology were calculated with the same linear regression model described for DMG identification.

#### 4. Pseudo-bulk compartment analysis

Pseudo-bulk chromatin contact matrices were generated for each cell type, donor, and brain region by merging raw single-cell contact files. Multi-resolution matrices were stored in.mcool format using cooler^54^. For each sample, 100 kb resolution contact matrices were used to compute eigenvectors of the pairwise contact matrices. PC1 values were sign-corrected based on bin GC content, such that A compartments had positive PC1 values and B compartments had negative PC1 values. These steps were performed with cooltools^55^, following the official online tutorial (https://cooltools.readthedocs.io/en/latest/notebooks/compartments_and_saddles.html).

The raw PC1 profiles from each sample and cell type were subjected to principal component analysis (PCA) to examine variation across samples. To identify genomic bins whose compartment status was associated with Alzheimer’s disease (AD) pathology, we applied the same linear regression framework used for differentially methylated genes (DMGs), as described above. Bins with an association P<0.05P < 0.05P<0.05 and an absolute regression coefficient |β|>0.25|\beta| > 0.25|β|>0.25 were considered to have differential PC1.

#### 5. Single-cell TAD analysis

TADs for each single cell were identified with scHiCluster, which implements the TopDom^56^ to call the TADs on an imputed 25 kb contact matrix. The association between the number of boundaries of each cell with AD pathology was calculated with the same linear regression model as described in DMG analysis. We quantified boundary activity in 1 Mb windows using binarized boundary calls at 25 kb resolution. For each cell, bins with any boundary signal were set to 1 (otherwise 0). We defined a sample as Region/Disease/Donor/Celltype and assigned each bin to a 1 Mb window using its genomic start coordinate.

For each sample × 1 Mb window, we counted the total number of boundary events by summing all (cell, bin) instances with value = 1 within that window, and divided by the total number of cells in the sample to obtain a per-cell normalized boundary density:

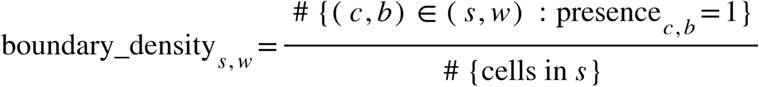

To mitigate coverage/quality differences, we computed the mean long-range cis contact per sample *mean CisLongContact* and regressed out this effect within each 1 Mb window *w* using an OLS model:

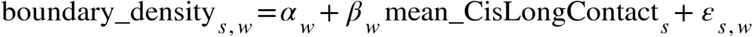

and took the residuals as the adjusted boundary density:

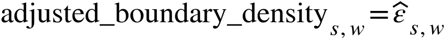

#### 6. Single-cell Loop analysis

The loops for each sample (Donor/Region/Celltype) were called with scHiCluster, following the online tutorial (https://zhoujt1994.github.io/scHiCluster/intro.html), which implements similar algorithms as SnapHiC^57^. We applied two steps to identify the differential loops:

a. Pairwise differential loops between sample groups For each chromosome in a given Cell/Region, pseudo-bulk contact matrices were generated separately for AD and Control groups by merging single-cell contact data from all samples in that group. Both the raw contact matrix and its element-wise squared matrix were loaded for each sample, multiplied by the number of contributing cells, and summed to produce group-level and overall cumulative matrices. Within-group variation (sum of squares) was calculated by iteratively subtracting the previous cumulative sum from the current group’s cumulative sum to obtain the group-specific totals, then combining these across the two groups. Total variation was computed from the overall cumulative matrices. The ratio of total to within-group variation was transformed into an F-statistic, providing a per-pixel measure of differential contact strength between AD and Control. To avoid artifactual signals, pixels overlapping blacklist regions (±7 bins) and those outside the genomic distance range of 5 to 500 bins were excluded. Only upper-triangle matrix entries were evaluated. The filtered per-pixel F-statistics were stored as sparse matrices, saved per chromosome for each Cell/Region, along with metadata including the total contributing cell count.
b. Linear regression model to identify loops associated with AD pathology Linear regression model described earlier in DMG identification was used to further regress out age and sex on the differential loops identified in step a.

### Opposing correlation between VC and PFC/TC with AD

Coefficients for each feature from DNA methylation, 3D genome (PC1, adjusted boundary density), and RNA data were obtained from the linear regression analyses described above, combining results across all cell types and brain regions. Only features significantly associated with Alzheimer’s disease (AD) pathology in at least one brain region were retained. Coefficients for these features were visualized using PyComplexHeatmap. Genes whose expression and CG DNA methylation exhibited concordant associations with AD across brain regions were selected for Gene Ontology (GO) enrichment analysis.

### scDRS analysis on Stressed and Homeostatic genes

We use the genes in the GO term “Alzheimer’s Disease” as representative genes associated with stress, while the genes in the GO term of “Neuron Development” as representatives of homeostatic genes. The gene module score for each set of genes were calculated with scDRS^22^, using the 10x multiome RNA data frome our companion study. We defined the RNA cell states with the stressed score and homeostatic score, using positive score 2.5, and negative score –2.5 as cutoff. Cells that are positive in both modules were assigned a double positive (DP) state, and both negatives were “DN”, others cell states were assigned according to the two scores. These RNA cell states were transferred to snm3C cells in the integration described above. The correlation between Stressed/Homeostatic states defined in DNA methylation and RNA cell states defined here were tested with fisher exact test.

### Putative telomere deletions analysis

#### 1. Per-sample centromere-aligned average contact (O/E)

For each sample, we loaded 100-kb cooler matrices (balance=True, ICE-normalized) at the chosen resolution. For each autosome with an available centromere position (excluding chrX/chrY and very small chromosomes with <50 bins), we:

a. **Centromere split.** Located the centromere bin by `searchsorted` on bin starts and split the chromosome contact map into four blocks: p–p, p–q, q–p, and q–q.
b. **Observed matrix normalization and resizing.** Took the observed matrix, replaced NaNs with 0, and resized each block to a fixed shape using bilinear interpolation (p_bins × p_bins for p–p, q_bins × q_bins for q–q, etc.). Reassembled the blocks into a standardized centromere-aligned matrix of size (p_bins+q_bins) × (p_bins+q_bins).
c. **Expected matrix from band means.** Estimated the distance-dependent expectation by computing the mean of each diagonal band of the observed matrix and reconstructing an “expected” matrix with those band means. Applied the same centromere split and resizing to build a standardized expected matrix.
d. **Per**-**chromosome O/E.** For each chromosome, calculate the observed-over-expected (O/E) centromere-aligned matrix.
e. **Average across chromosomes.** Averaged the standardized O/E matrices over all eligible chromosomes to obtain one centromere-aligned average O/E contact matrix per sample. Any NaN/Inf values were set to 0.
f. The resulting per-sample matrices summarize arm-scaled, centromere-aligned contact patterns after removing distance-decay effects via the expected (band-mean) model.

The averaged contact matrices across all chromosomes for each cell type/brain region/Cell states were plotted.

#### 2. Detection of bins with widespread contact loss (putative deletions)

For each brain region (VC, PFC, TC) and major cell type, we generated state-specific pseudo-bulk Hi-C maps (Homeostatic, Stressed) and loaded ICE-balanced contact matrices at 100 kb resolution from.mcool files. For every autosome, we computed the row sum of the observed, balanced contact matrix (sum across columns per bin), yielding a per-bin contact intensity for each (Region × Cell type × State) pseudo-bulk.

Per chromosome, these row-sum vectors were assembled into a data frame with bins as rows and pseudo-bulks as columns (columns renamed to State_CellType_Region). To robustly flag bins showing near-absence of contacts, zeros were recoded to NaN, and bins with ≥ 10 missing values across pseudo-bulks were selected. After sorting columns and removing rows that were entirely missing, the remaining bins constituted the set of putative deletions on that chromosome. Results from all chromosomes were concatenated to produce the final putative deletions. The distance of these deletions was calculated and GO enrichment of genes in these regions was examined using gprofiler.

### Epigenomic analysis of Homeostatic and Stressed cell states

#### 1. Compartment analysis of different cell states across cell types

Pseudo-bulk Hi-C contact matrices for the Homeostatic and Stressed cell states in each cell type and brain region were generated by merging single-cell contact files. PC1 values for each pseudo-bulk were computed as described above for the AD versus control comparison. Saddle plots were generated with *cooltools* following the same online tutorial referenced previously. To assess compartment shifts, the observed/expected saddle plot matrix for the Stressed state was divided by the corresponding Homeostatic matrix, and the resulting fold-change values were visualized.

#### 2. Boundary density and TAD-centered pileup plots

Adjusted boundary densities for the two cell states were calculated using the same approach as for the AD versus control analysis described above. Boundary density profiles were plotted with *PyComplexHeatmap*, clustering columns separately for Homeostatic and Stressed states. TAD-centered pileup plots were computed with *coolpuppy* using parameters ––local ––rescale_flank 1 ––rescale ––rescale_size 99. Pileup matrices for the Stressed and Homeostatic states were then subtracted, and the differences were visualized with *plotpup.py*.

#### 3. Differential loops between the two cell states

Differential loops between Homeostatic and Stressed states were identified using the same statistical framework as for the AD versus control comparisons across brain regions and cell types.

#### 4. Differentially expressed genes between the two cell states

DEGs between the two cell states in each brain region were also identified with MAST in Seurat, regressing out “nFeature_RNA”, “Age”, “Sex”, “Region”, “Disease”.

## Data Availability

All snm3C sequencing data generated in this study have been deposited in the NCBI Gene Expression Omnibus (GEO) under accession number **GSE308132**. The sample level methylation data can be explored at http://neomorph.salk.edu/aj2/pages/hchen/human_ad3c.php. The 10x Multiome datasets generated in the companion study will be made available upon their submission to GEO.

## Code Availability

All computational tools used in this study are publicly available. Software developed by others, as well as any online tutorials followed, are appropriately cited in the Methods section. Custom scripts for data processing and analysis are available at the project’s GitHub repository: https://github.com/wangwl/ADsnm3C.

## Author Contributions

Conceptualization: J.R.E, B.T.H, Methodology: W.W., Sample Processing: ADRC team at MGH, Nuclei Sorting: M.L., M.M., C.O., Data Generation: R.C., A.B., J.R.N.,M.K., C.V, J.A., A.P., D.C., C.C., A.S.A., J.L., M.J., E.S., Investigation: W.W., Formal analysis: W.W., P.B., B.Y., Writing – original draft: W.W., Writing – review & editing: All authors, Discussion: N.Z., D.O., Supervision: B.R., B.T.H., J.R.E. Funding acquisition: B.R., B.T.H, J.R.E..

## Acknowledgments

This work was supported by grants from National Institute on Aging (NIA) 7R56AG069107-02 to B.R., B.T.H. and The American Heart Association (AHA) 611612 to J.R.E.. J.R.E. is an Investigator of the Howard Hughes Medical Institute. All human brain tissues and fibroblasts were obtained from the Massachusetts Alzheimer’s Disease Research Center (MADRC, P30AG062421). We would like to thank all the unidentified donors who contributed biological samples for this project through our collaborators. K.K. was supported by the David Goeddel Fellowship and the National Science Foundation Graduate Research Fellowship Program under Grant No. DGE-2038238. Any opinions, findings, and conclusions or recommendations expressed in this material are those of the author(s) and do not necessarily reflect the views of the National Science Foundation. We are thankful to the FACS core for their help in processing the samples. This work utilized the Stampede2 supercomputing resources at Texas Advanced Computing Center (TACC) through the Extreme Science and Engineering Discovery Environment (XSEDE) and the Anvil supercomputing resources at Purdue University’s Rosen Center for Advanced Computing (RCAC) made available through the Advanced Cyberinfrastructure Coordination Ecosystem: Services & Support (ACCESS) program funded by the National Science Foundation (NSF), grant numbers 1540931 and 2005632 respectively, via Research Allocation to the Ecker lab grant MCB130189.

## Declaration of Interests

J.R.E. is a scientific advisor for Zymo Research Inc. and Ionis Pharmaceuticals. B.R. is a co-founder and consultant of Arima Genomics and co-founder of Epigenome Technologies.

**Extended Data Figure 1.**
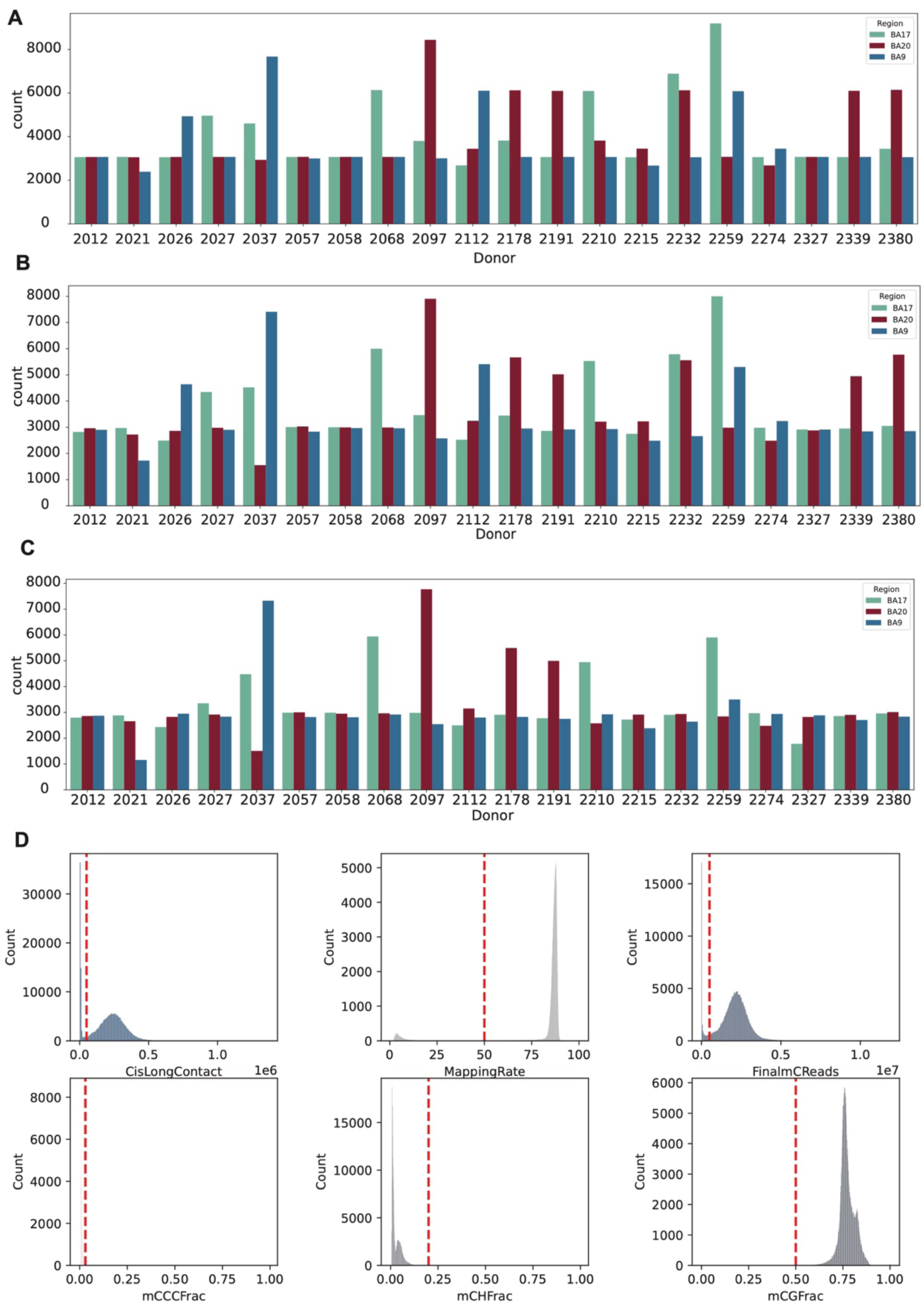
Number of cells sequenced and retained after filtering, and filtering criteria. (A) Number of cells sequenced for each donor in three brain regions. (B) Number of cells passing DNA methylation QC for each donor in three brain regions. (C) Number of cells passing both DNA methylation and 3C QC for each donor in three brain regions. (D) Metrics used for cell filtering and the distribution of these metrics across all cells.

**Extended Data Figure 2.**
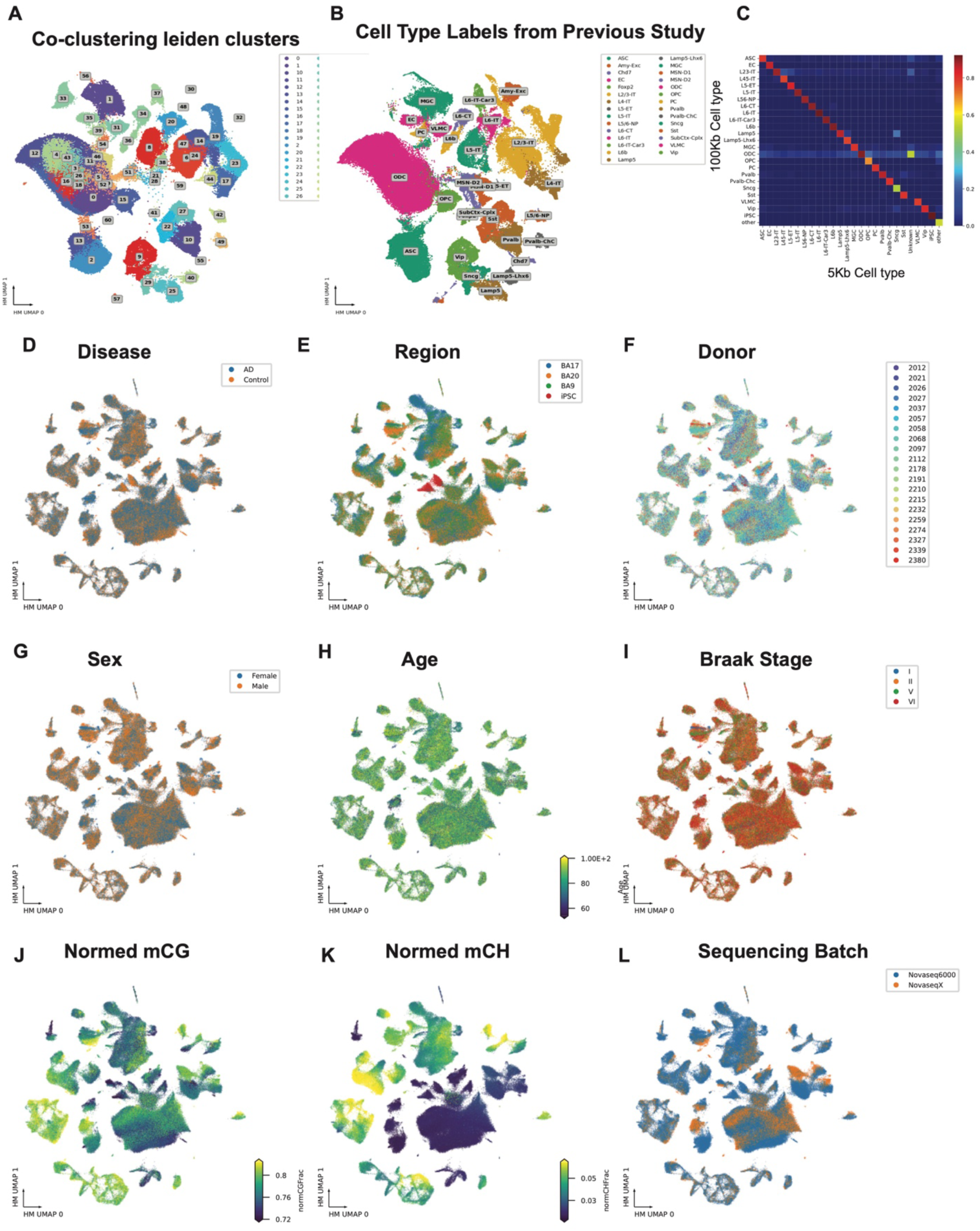
UMAPs of clustering results. (A) Co-clustering Leiden clusters of snm3C cells from this study, colored by Leiden cluster. (B) Co-clustering of cells from a previous study, colored by cell types annotated in that study. (C) Heatmap showing the ratio of cells annotated at 5 kb resolution that are assigned to cell types at 100 kb resolution. (D–L) UMAPs of 5 kb CG methylation clustering results, colored by different metadata as indicated above each panel.

**Extended Data Figure 3.**
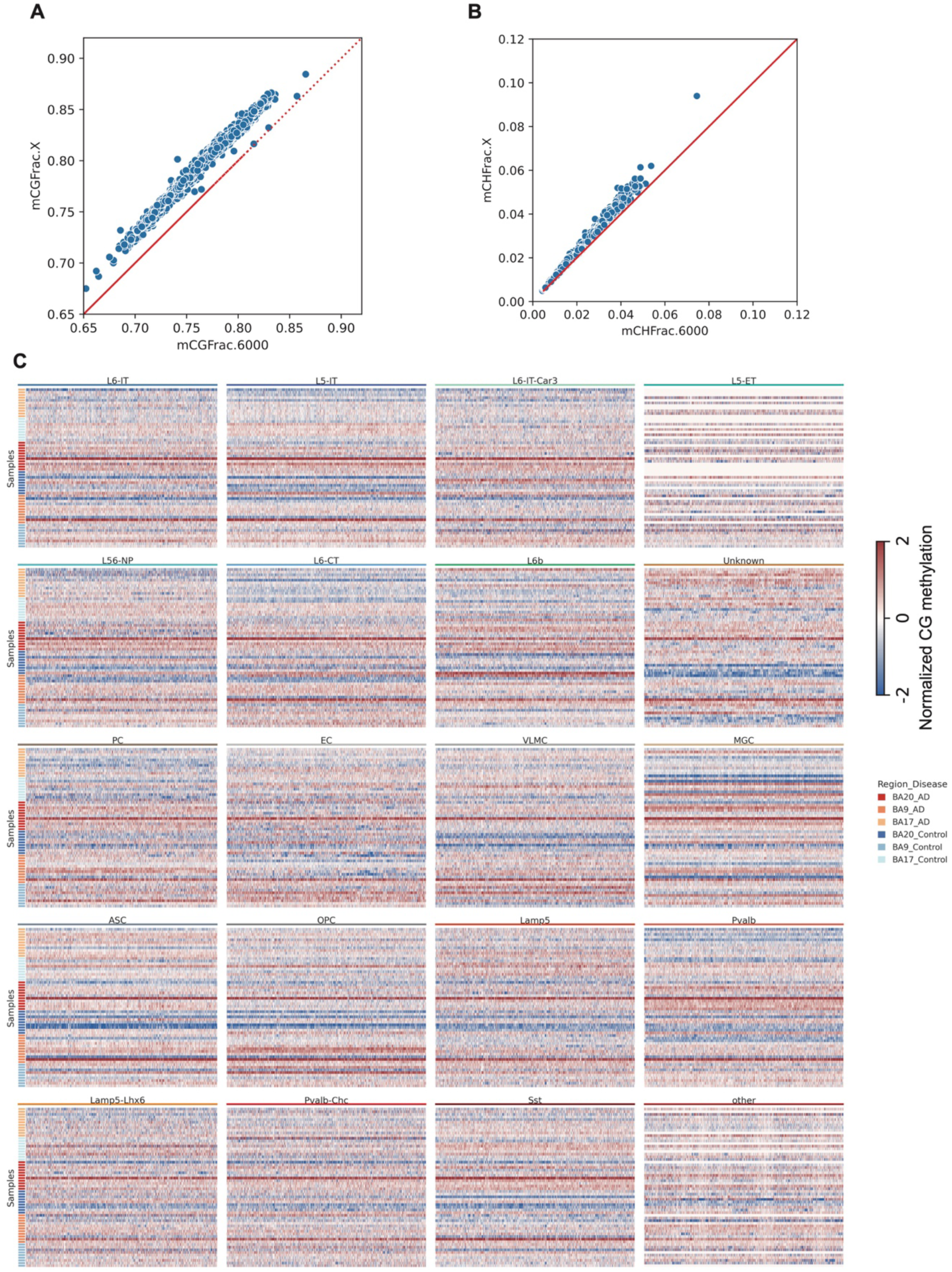
Genome-wide normalized CG methylation levels at 1 Mb resolution. Genome-wide normalized CG methylation levels at 1 Mb resolution across brain regions in AD and control samples for all cell types.

**Extended Data Figure 4.**
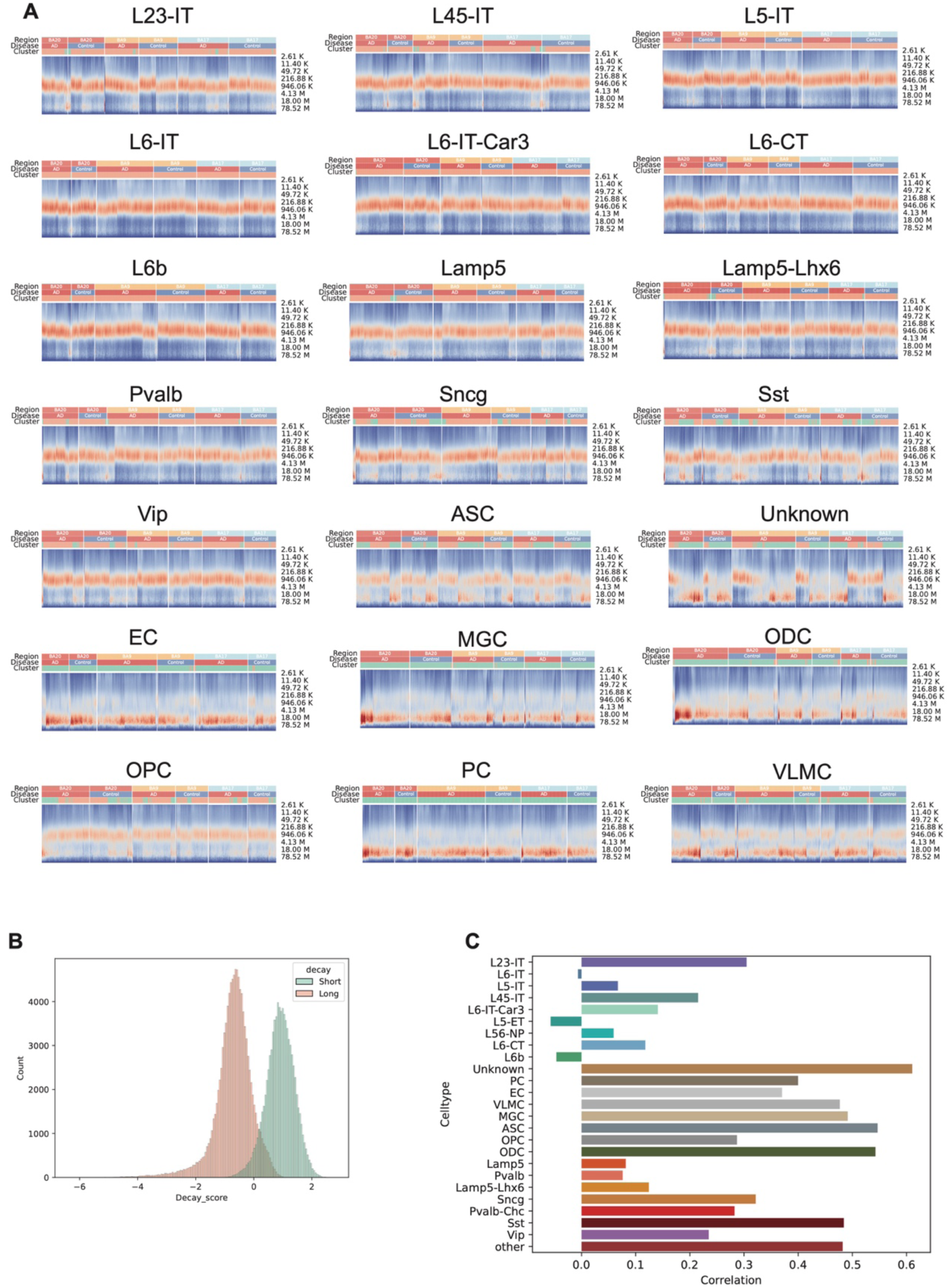
Number of contacts at different genomic distances. **(A)** Heatmaps show the relationship between contact frequency and genomic distance in single cells across all cell types. Colors indicate the fraction of contacts at each distance; annotations indicate brain region and disease state. Cells enriched in long-range contacts are classified as Long, and others as Short. **(B)** Histogram shows the distribution of contact scores in cells classified as short contact–enriched or long contact–enriched. **(C)** Correlation coefficients between contact score and number of boundaries in each single cell across all cell types.

**Extended Data Figure 5.**
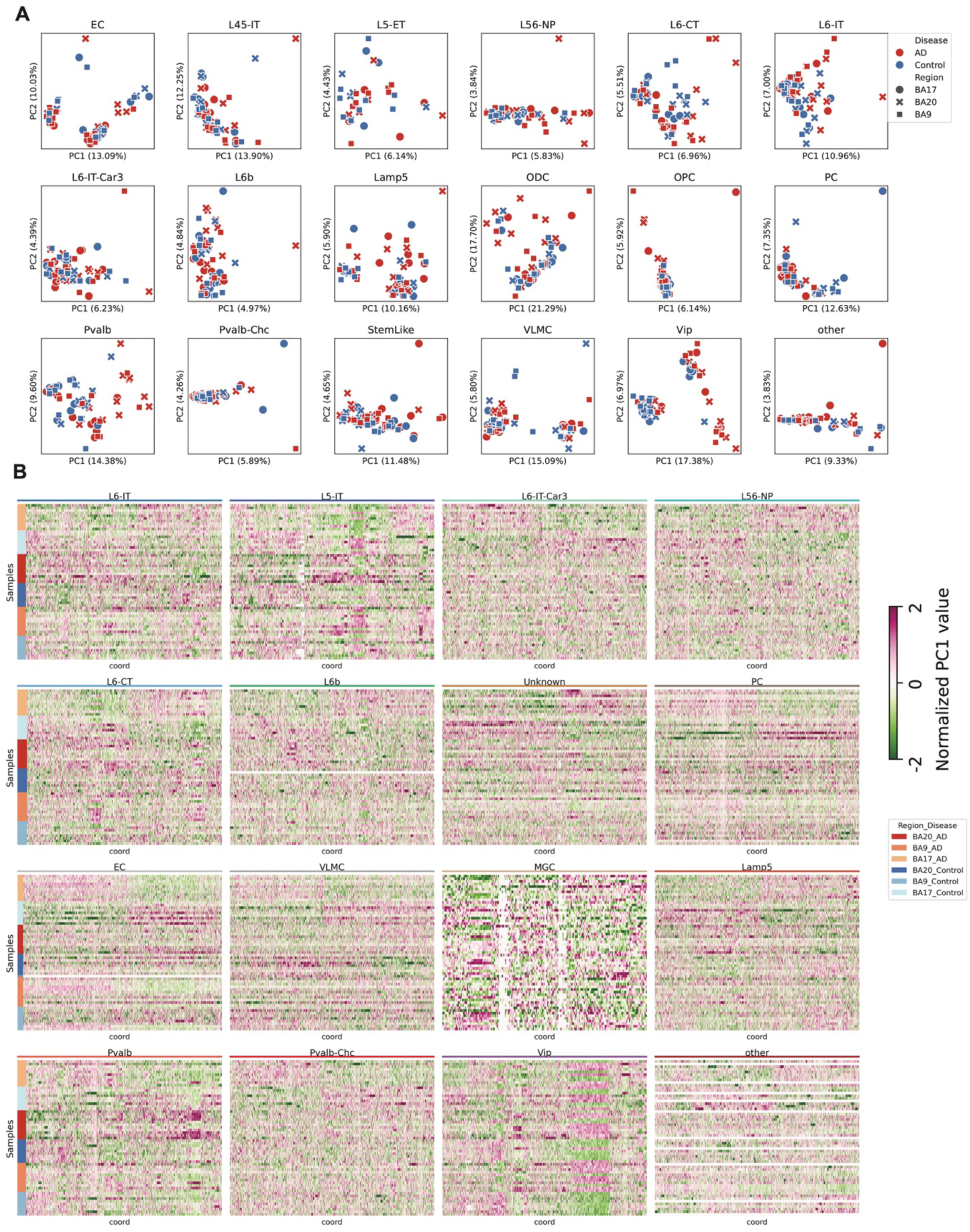
Compartment analysis between AD and controls in all cell types and brain regions. (A) Scatter plots show the principal component analysis (PCA) of PC1 values from 3D genome compartment profiles. (B) Heatmaps display the normalized PC1 values across samples for genomic bins whose PC1 values are significantly associated with AD, either positively or negatively.

**Extended Data Figure 6.**
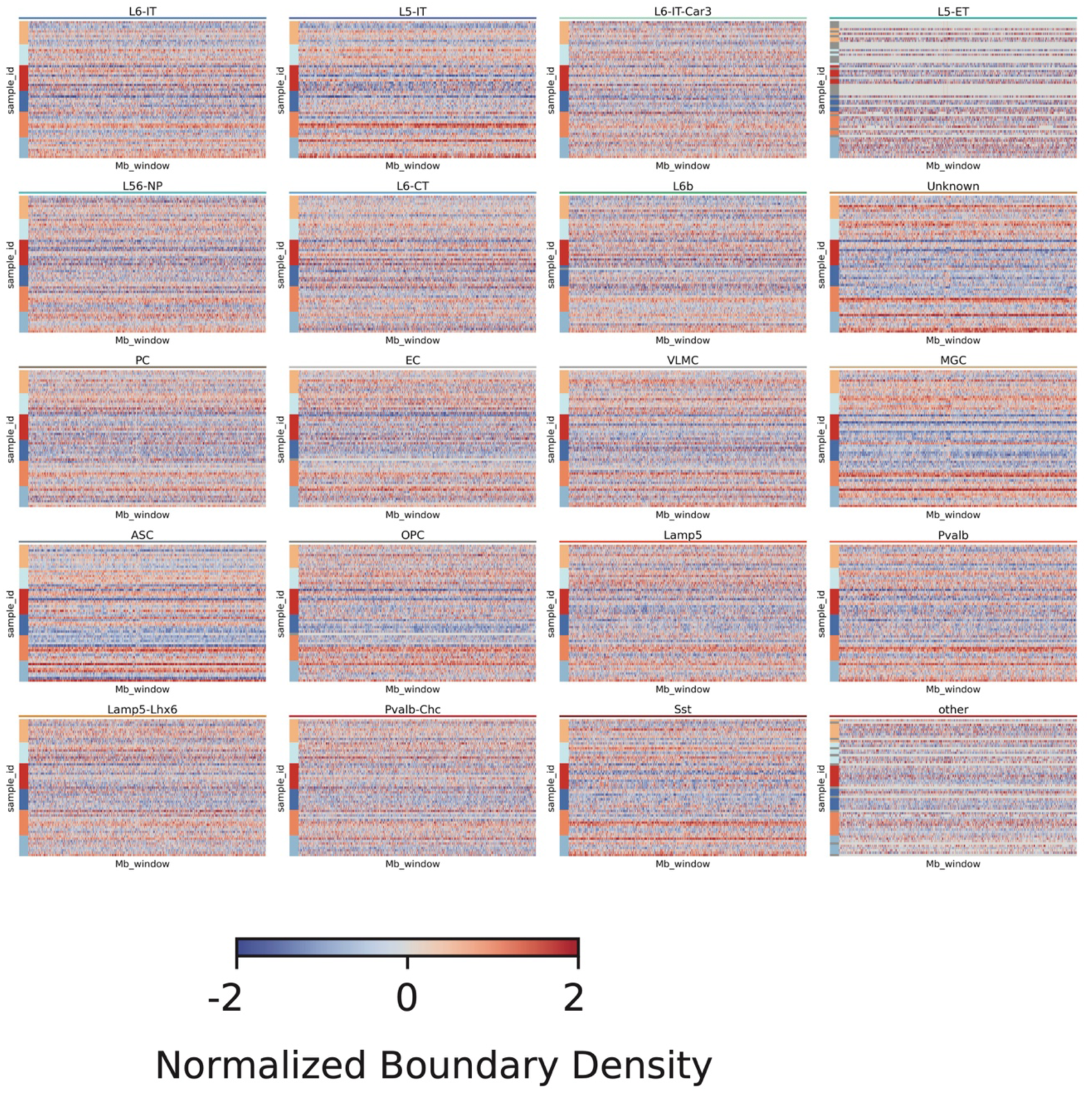
Normalized boundary densities in AD and controls across all cell types and brain regions. Heatmaps show the normalized boundary densities in AD and control samples across all cell types and brain regions.

**Extended Data Figure 7.**
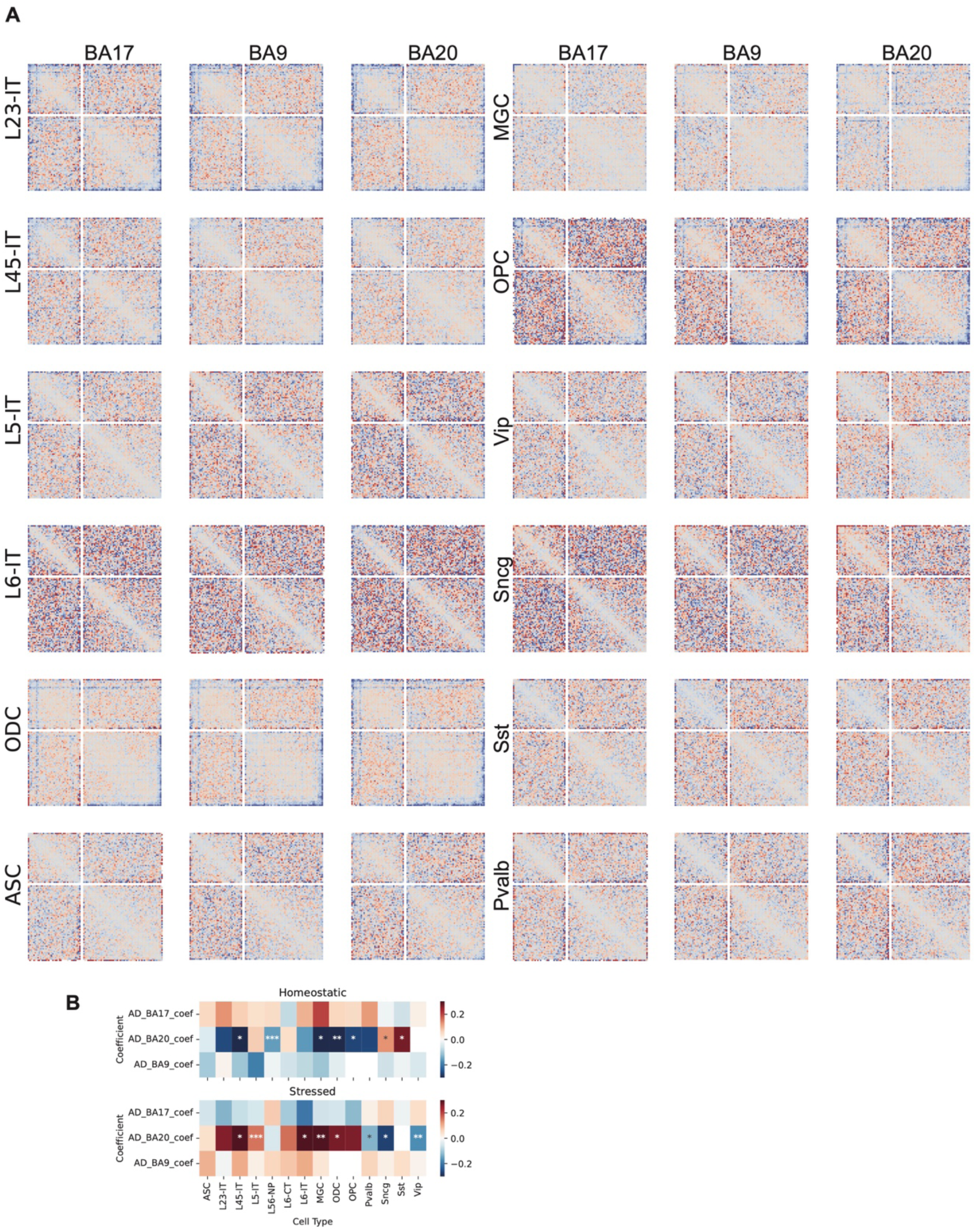
Differential global contacts between stressed and homeostatic states. Heatmaps show the division of averaged obs/exp contact maps across all chromosomes, with scaled p– and q-arms, between stressed and homeostatic states across three brain regions and all cell types.

**Extended Data Figure 8.**
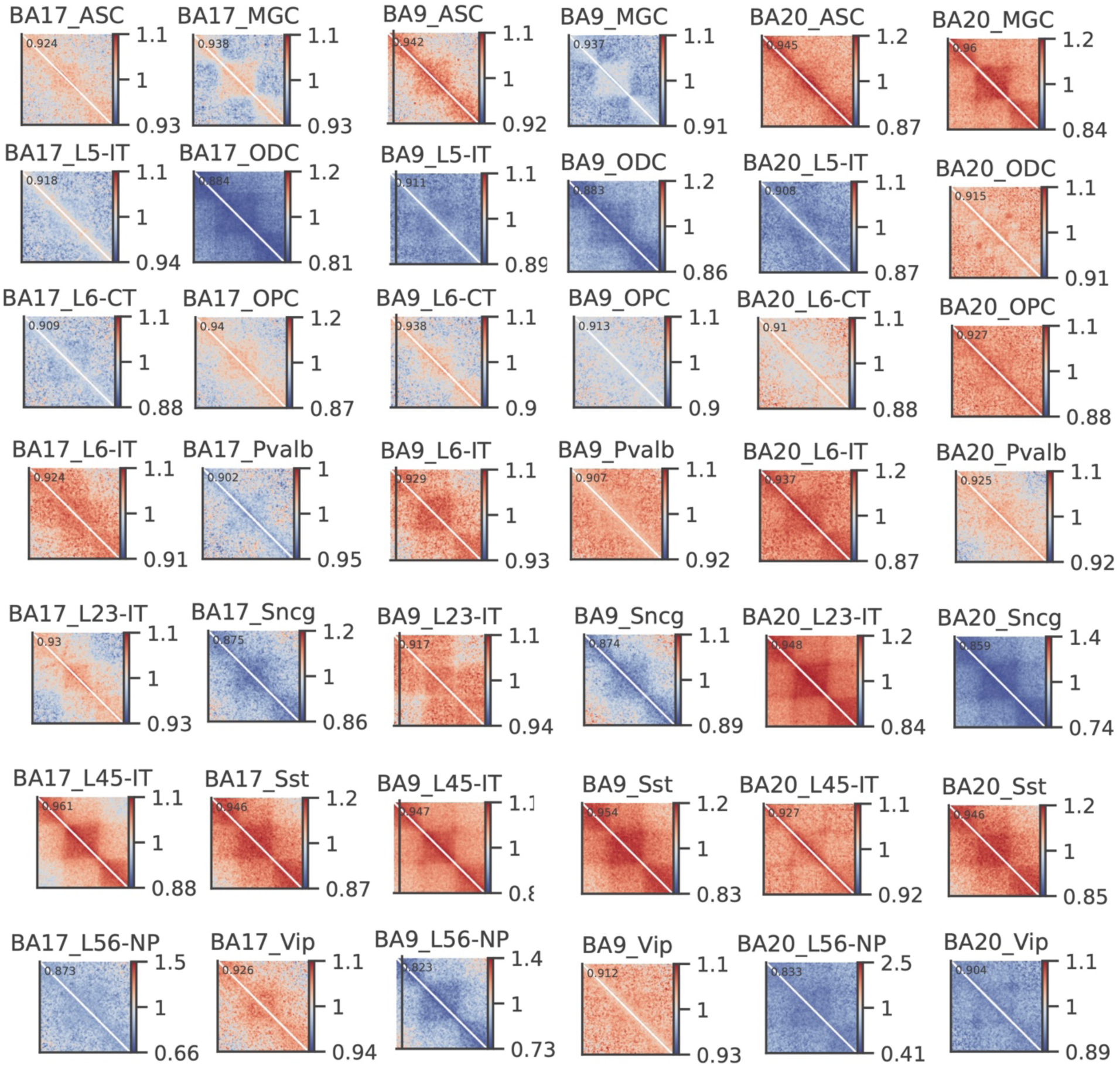
Difference of short-range contacts between stressed and homeostatic states. Division (stressed / homeostatic) of pileup matrices centered on TADs across all cell types and three brain regions.

